# Transferrin plays a central role to maintain coagulation balance by interacting with clotting factors

**DOI:** 10.1101/646075

**Authors:** Xiaopeng Tang, Zhiye Zhang, Mingqian Fang, Yajun Han, Sheng Wang, Min Xue, Yaxiong Li, Li Zhang, Jian Wu, Biqing Yang, Qiumin Lu, Xiaoping Du, Ren Lai

## Abstract

Coagulation balance is maintained through fine-tuning interactions among clotting factors. Physiological concentrations of clotting factors are huge difference. Especially, coagulation proteases’ concentration (pM to nM) is much lower than their natural inactivator antithrombin III (AT-III, ∼3 μM). Here we show that transferrin (normal plasma concentration ∼40 μM) interacts with fibrinogen, thrombin, FXIIa and AT-III with different affinity to maintain coagulation balance. Normally, transferrin is sequestered by binding with fibrinogen (normal plasma concentration ∼10 μM) with a molar ratio of 4:1. In atherosclerosis, abnormally up-regulated transferrin interacts with and potentiates thrombin/FXIIa and blocks AT-III’s inactivation on coagulation proteases by binding to AT-III, and thus induces hypercoagulability. In mouse models, transferrin-overexpression aggravated atherosclerosis while transferrin-knockdown, anti-transferrin antibody or designed peptides interfering transferrin-thrombin/FXIIa interactions alleviated it. Collectively, these findings identify transferrin as a clotting factor and an adjuster for maintaining coagulation balance and modify the coagulation cascade.

## Introduction

Normal coagulation pathway represents a balance between pro-coagulant and anti-coagulant pathway through fine-tuning interactions among clotting factors(Adams and Bird, 2009; Davie et al., 1991; Esmon, 1989, 2000; Palta et al., 2014; Smith et al., 2015). Nineteen clotting factors have been identified to take part in the coagulation cascade and to maintain the thrombohemorrhagic balance (Esmon, 2000; Jesty and Beltrami, 2005; Palta et al., 2014; Perez-Gomez and Bover, 2007). They include fibrinogen family (fibrinogen (factor I), factor V, factor VIII and factor XIII) (Triplett, 2000), vitamin K dependent family ((factor II, thrombin), factor VII, factor IX and factor X) (Bhattacharyya et al., 2005; Mann et al., 1990), contact family (factor XI, factor XII, high molecular weight kininogen (HMWK, factor XV) and prekallikerin (PK, factor XIV)) (Schmaier, 2016; Travers et al., 2015), tissue factor (factor III), calcium (factor IV), Von Willebrand factor (vWf, factor XVI), antithrombin III (AT-III, factor XVII), heparin cofactor-II (factor XVIII), protein C (factor XIX) and protein S (factor XX) (Davie, 2003; Newland, 1987; Palta et al., 2014; Periayah et al., 2017; Schmaier, 2016). They are resembled a coagulation waterfall or cascade comprising of the intrinsic and the extrinsic pathways, which converge in FX activation (Davie, 2003; Hoffman, 2003; Hougie, 2004). In this model, each clotting factor consists of a proenzyme that is converted to an active enzyme by the upstream activated clotting factor (Furie and Furie, 1988). Normally, the coagulation process is under the inhibitory control of several inhibitors to create a physiological balance between anticoagulant and antifibrinolytic pathways, which is named coagulation balance to avoid too much and too little clot production (Chan and Paredes, 2013; Esmon, 1989; Ganter and Spahn, 2010; Mann et al., 2009; Palta et al., 2014; Roderique and Wynands, 1967).

Fine-tuning interactions among clotting factors play key roles for maintaining coagulation balance but the concentration of clotting factors is huge difference (Palta et al., 2014). Normally, the concentration of fibrinogen, prothrombin, and AT-III is ∼10, ∼2 and ∼3 μM, respectively, while that for most of coagulation proteases is pM to nM (Esmon, 2000). AT-III is the main natural inactivator of coagulation proteases to limit the clot formation, thus avoiding the thrombus propagation (Esmon, 2000). Given the huge concentration difference of different clotting factors, especially that the concentration of coagulation proteases is thousands times lower that of their physiological inactivator AT-III, we suppose that there are some adjusters or balancers to orchestrate or buffer imbalanced clotting factors with different concentration, or to sequester interactive objects in normal plasma. We found that transferrin (Tf), an endogenous plasma protein with high concentration of 40 μM that transports iron (Gkouvatsos et al., 2012; Prinsen et al., 2001), acts as a clotting factor and an adjuster for maintaining coagulation balance by interacting with not only clotting factors with high concentration, i.e., fibrinogen and AT-III, but also clotting factors with low concentration, i.e., coagulation proteases. Usually, most of Tf is sequestered from coagulation proteases and AT-III by binding with fibrinogen.

## Results

### Enhanced enzymatic activity of thrombin and FXIIa is associated with elevated Tf in atherosclerosis

In order to investigate the mechanism responsible for atherosclerosis (AS)-associated hypercoagulability, enzymatic activities of several coagulation factors including FVIIa, FXIa, FXIIa, kallikrein and thrombin were compared between plasma from healthy people and atherosclerotic patients (coronary heart disease (CHD) patients) who had angiographically-visible luminal narrowing (Table S1). The enzymatic activity of thrombin and FXIIa in atherosclerotic plasma was found to be 2-2.5 times stronger than that in healthy plasma (Figure S1A), although total levels of prothrombin and FXII are not different (Figures S1B-S1D), suggesting the presence of factors potentiating the activation or activities of thrombin/FXIIa in atherosclerotic plasma. The enzymatic activity of kallikrein, FXIa and FVIIa are not significantly different between plasmas from AS patients and normal controls (Figure S1A). The factor with potentiating effects on thrombin/FXIIa was purified and identified as Tf (Figures S2). An anti-Tf antibody (Tf AB) blocked the elevation of the enzymatic activities of thrombin (Figure 1A) and FXIIa (Figure 1B) in the atherosclerotic plasma, further suggesting that Tf is a potentiator for these two coagulation factors. Comparative analysis using enzyme linked immunosorbent assay (ELISA) and western blot confirmed that there are elevated Tf in atherosclerotic plasma (Figures S1C and S1D). The average Tf concentration in the plasma of CHD patients (n = 42, male 22; female 20) was 4.173 mg/ml (SD 1.19), while that in healthy individuals (n = 42, male 21; female 21) was 2.865 mg/ml (SD 0.33) (Figure 1C). High levels of Tf were also observed in the atherosclerotic lesion specimens (Figure 1E and Table S2).

**Figure 1.**
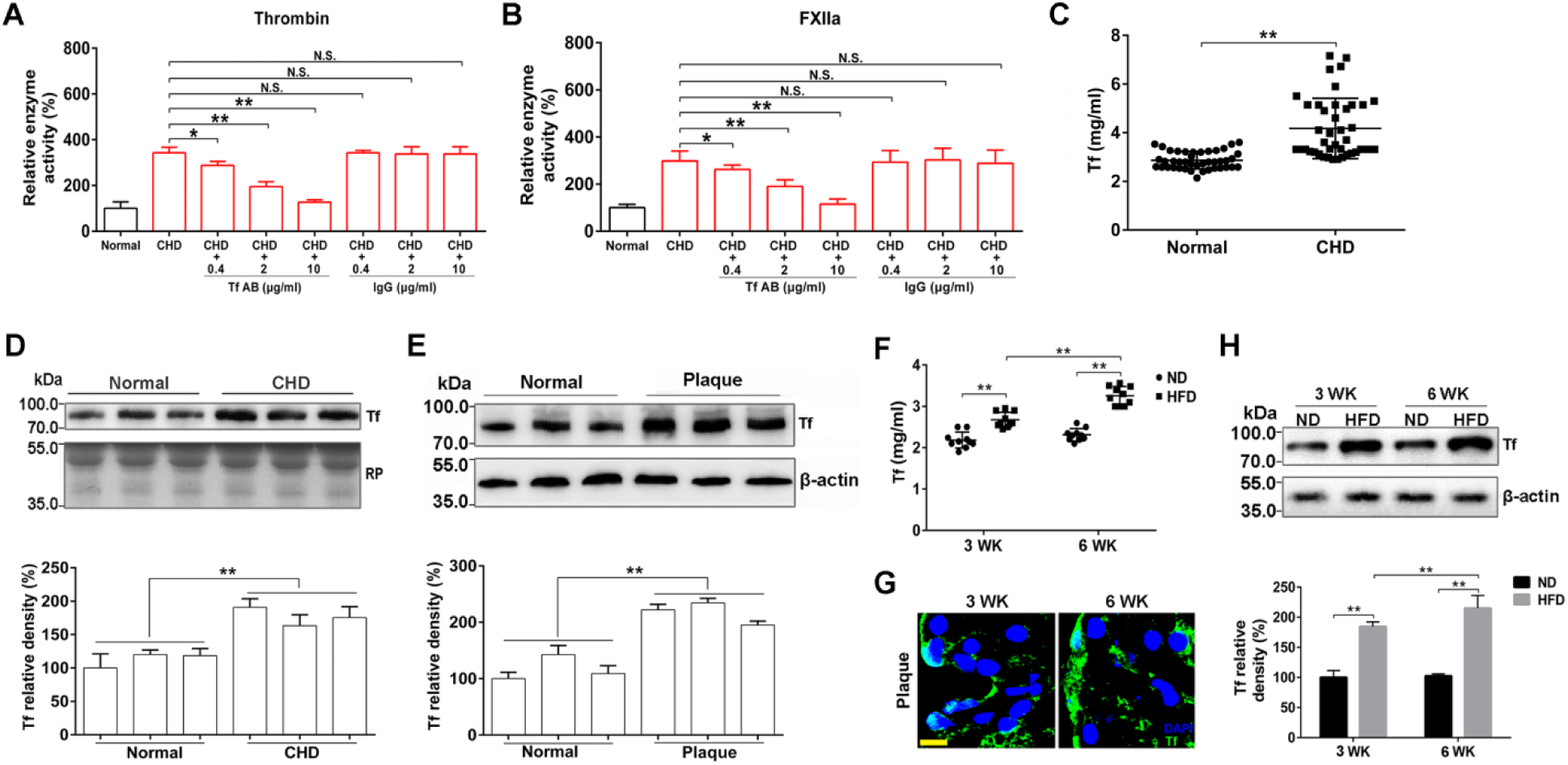
Enhanced enzymatic activity of thrombin and FXIIa is associated with elevated Tf in atherosclerotic plasma. An anti-Tf antibody (Tf AB) alleviated the potentiating ability of CHD plasma on enzymatic activity of thrombin **(A)** and FXIIa **(B)**. Data represent mean ± SD (n= 6), \*\**p* < 0.01 by one-way ANOVA with Dunnett’s post hoc test. **(C)** Amounts of Tf in plasma from CHD patients and volunteers were determined by ELISA. Data represent mean ± SD (n = 42), \*\**p* < 0.01 by unpaired t-test. **(D)** Western blot (top) and quantification (bottom) analysis of Tf in plasma samples from CHD patients and health volunteers. Red Ponceau (RP)-stained blot is used as a loading control. **(E)** Western blot (top) and quantification (bottom) analysis of protein extracts from normal arteries (Normal) and atherosclerotic lesions (Plaque). β-actin is used as the control. Data represent mean ± SD (n = 12), \*\**p* < 0.01 by unpaired t-test **(D, E). (F)** Amounts of Tf in the plasma from the *Apoe*^−*/*−^ mice fed with high fat diet (HFD)- and normal diet (ND)-fed mice were determined by ELISA. Data represent mean ± SD (n = 10), \*\**p* < 0.01 by unpaired t-test. **(G)** Immunofluorescence staining of Tf (green) in mice atherosclerotic plaque. Cell nuclei were labeled by DAPI. Scale bar represents 10 μm. Images were representative of at least three independent experiments. **(H)** Western blot (top) and quantification (bottom) analysis of Tf in aortic roots of the *Apoe*^−*/*−^ mice. Data represent mean ± SD (n = 10), \*\**p* < 0.01 by unpaired t-test. N.S.: no significance.

To further investigate the association of elevated Tf with atherosclerosis, the *Apoe*^−*/*−^ mice were fed a normal diet (ND) or a high fat diet (HFD, 21% fat, 0.15% cholesterol) for 6 weeks to test the change of Tf in the plasma and atherosclerotic plaque. Notably, elevated Tf level was observed in the plasma of HFD fed *Apoe*^−*/*−^ mice (Figure 1F), which was congruent with atherosclerotic plaque development (Figure S3A). Confocal microscopy and immunoblot analysis also showed increased Tf in atherosclerotic plaque (Figures 1G and 1H). In addition, quantitative real-time polymerase chain reaction (qRT-PCR) showed the *Tf* RNA is dominantly up-regulated in liver, indicating that liver is the main site of Tf synthesis (Figure S3B).

### Tf potentiates thrombin/FXIIa and inhibits AT-III independently of iron

As an iron carrier, Tf exists in plasma in both the ferric iron-bound state (holo-Tf) or unbound state (apo-Tf). As illustrated in Figures 2A and 2D, both apo-Tf and holo-Tf were found to show a similar effect to enhance the enzymatic activities of thrombin and FXIIa. At the concentration of 0.2, 1 and 5 μM, Tf enhanced the enzymatic activity of thrombin by ∼0.2, 1 and 1.8-fold, while that for FXIIa was ∼0.2, 0.7 and 1.5-fold, respectively. Similarly, apo-Tf and holo-Tf exhibited no difference in promoting coagulation through shortening the recalcification time (Figure S4). The effects of Tf on thrombin and FXIIa were further demonstrated by enhancing the enzymes’ ability to hydrolyze their natural substrates, fibrinogen (Figures 2B and 2C) and prekallikrein (PK) (Figures 2E and 2F), respectively. Fibrinopeptide A (FbpA) resulted from fibrinogen hydrolysis by thrombin was increased ∼0.2, 0.5, and 1.2-fold by 0.2, 1 and 5 μM Tf in 30 min, while that for FbpB was ∼1.1, 2.1, and 4.2-fold, respectively (Figures 2B and 2C). At the concentration of 0.2, 1 and 5 μM, Tf increased the ability of FXIIa to release the hydrolytic product (kallikrein heavy chain (HC), 52 kDa) of PK by 0.8, 1.9 and 2.7-fold, respectively (Figures 2E and 2F). Tf showed effects on neither zymogen activation of thrombin and FXIIa nor activities of kallikrein, FXIa and FVIIa (Figure S5). As illustrated in Figures 2G and 2I, both apo-Tf and holo-Tf blocked the inhibitory activity of AT-III towards thrombin and FXa. The inactivation on thrombin and FXa by 2 μM AT-III was completed blocked by 10 μM Tf. As a result, generation of thrombin-AT-III (TAT) and FXa-AT-III complexes were blocked (Figures 2H and 2J). In addition, thrombin-induced platelet aggregation was augmented by Tf (Figure S6). These data indicate that Tf induces hypercoagulability by potentiating thrombin and FXIIa and blocking inactivation of antithrombin III on thrombin and FXa.

**Figure 2.**
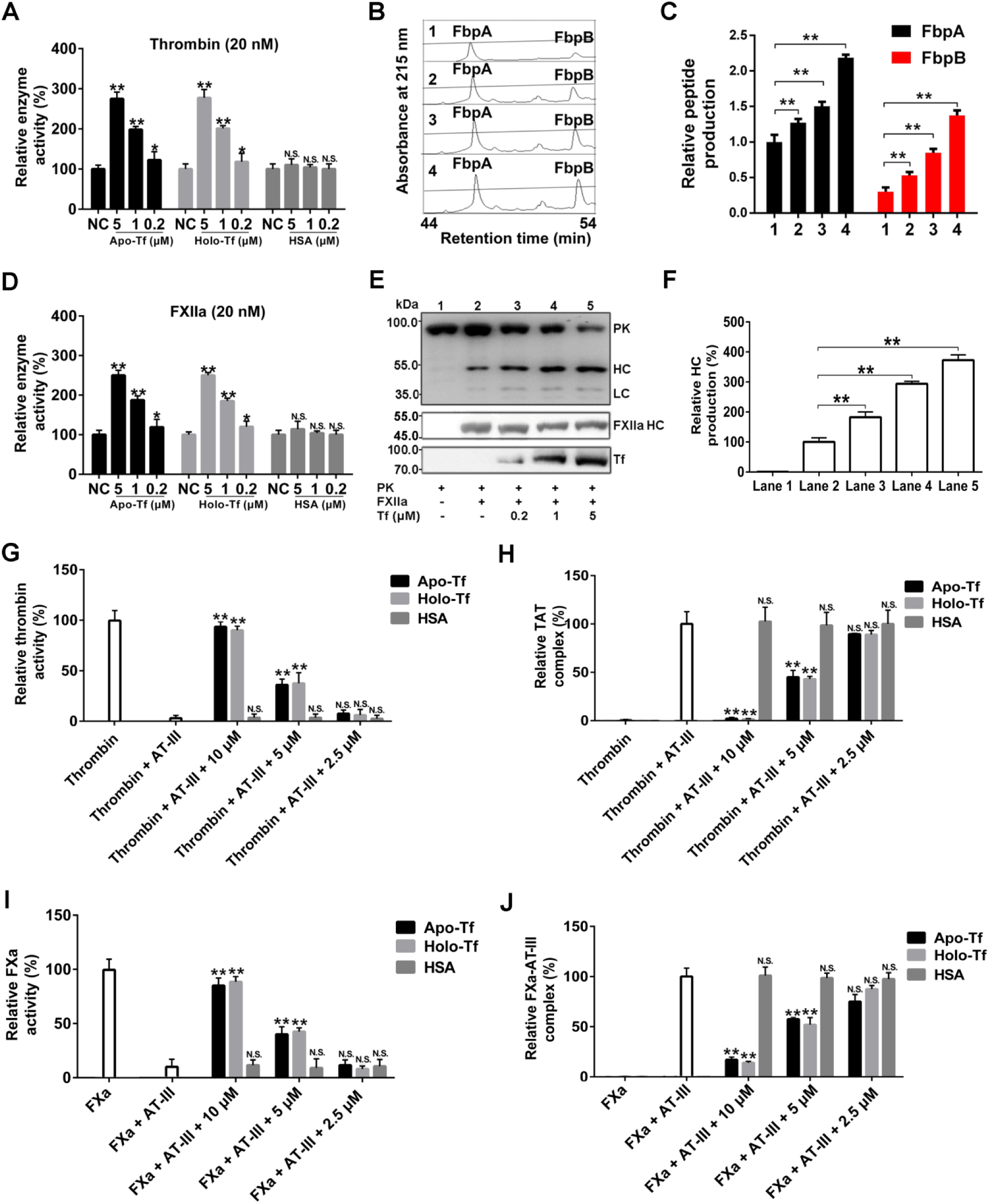
Effects of both apo- and holo-Tf on thrombin, FXIIa and antithrombin. **(A)** The potentiating effects of both apo- and holo-Tf on thrombin. (**B)** Representative RP-HPLC analysis and quantification **(C)** of fibrinopeptide A (FbpA) and fibrinopeptide B (FbpB) released from 5 mg fibrinogen hydrolyzed by 0.1 NIH unit thrombin mixed with 0, 0.2, 1 and 5 μM apo-Tf (panel 1-4), respectively. **(D)** The potentiating effects of both apo- and holo-Tf on FXIIa. Representative western blot **(E)** and quantification analysis of kallikrein heavy chain (HC∼52 kDa) **(F)** released from 10 μg prekallikrein (PK) hydrolyzed by 0.01 NIH unit FXIIa mixed with 0, 0.2, 1 and 5 μM apo-Tf (lane 2-5), respectively. Blotting of PK, FXIIa heavy chain (FXIIa HC), Tf and kallikrein light chain (LC∼36 and 33 kDa) were also shown. Apo- and holo-Tf block antithrombin III (AT-III)’s inactivation on thrombin **(G, H)** and FXa **(I, J)**. TAT: thrombin-AT-III complex. HSA: human serum albumin. Data represent mean ± SD of five independent experiments, \**p* < 0.05, \*\**p* < 0.01 by one-way ANOVA with Dunnett’s post hoc test. N.S.: no significance.

### Tf directly interacts with thrombin, FXIIa, fibrinogen and AT-III

Surface plasmon resonance (SPR) analysis revealed that apo-Tf directly interacts with thrombin (Figure 3A), FXIIa (Figure 3B), fibrinogen (Figure 3C) and AT-III (Figure 3D), whereas human serum albumin (HSA, negative control) shows no interaction with them (Figure S5D). The binding molar ratio between apo-Tf and fibrinogen or AT-III is 4:1 or 2:1 (Figure S5E). The association (*Ka*), dissociation (*Kd*) rate constants and equilibrium dissociation constant (*KD*) of the interaction between apo-Tf and thrombin was 4.7 × 10^5^ M^−1^s^−1^, 3.6 × 10^−3^ s^−1^ and 7.7 nM, while that for apo-Tf-FXIIa interaction was 1.8 × 10^5^ M^−1^s^−1^, 2.5 × 10^−3^s^−1^ and 13.9 nM respectively. The parameter was 3.4 × 10^4^ M^−1^s^−1^, 1 × 10^−3^ s^−1^ and 29 nM for apo-Tf-fibrinogen interaction, and 2.1 × 10^3^ M^−1^s^−1^, 1.1 × 10^−3^ s^−1^ and 524 nM for apo-Tf-AT-III interaction, respectively. The native gel shift assays also showed a complex formation between apo-Tf and thrombin (Figure 3E), FXIIa (Figure 3F), fibrinogen (Figure 3G) or AT-III (Figure 3H). SPR analysis also revealed that apo-Tf interacted with prothrombin and FXII (Figures S5F and S5G). *Ka, Kd* and *KD* for apo-Tf-prothrombin interaction was 1.4 × 10^5^ M^−1^s^−1^, 2.5 × 10^−3^ s^−1^ and 18 nM, and 0.3 × 10^5^ M^−1^s^−1^, 1.2 × 10^−3^s^−1^ and 40 nM for apo-Tf-FXII interaction, respectively. Holo-Tf also interacted with thrombin, FXIIa, fibrinogen, AT-III, prothrombin and FXII with the similar property of apo-Tf (Figure S5H).

**Figure 3.**
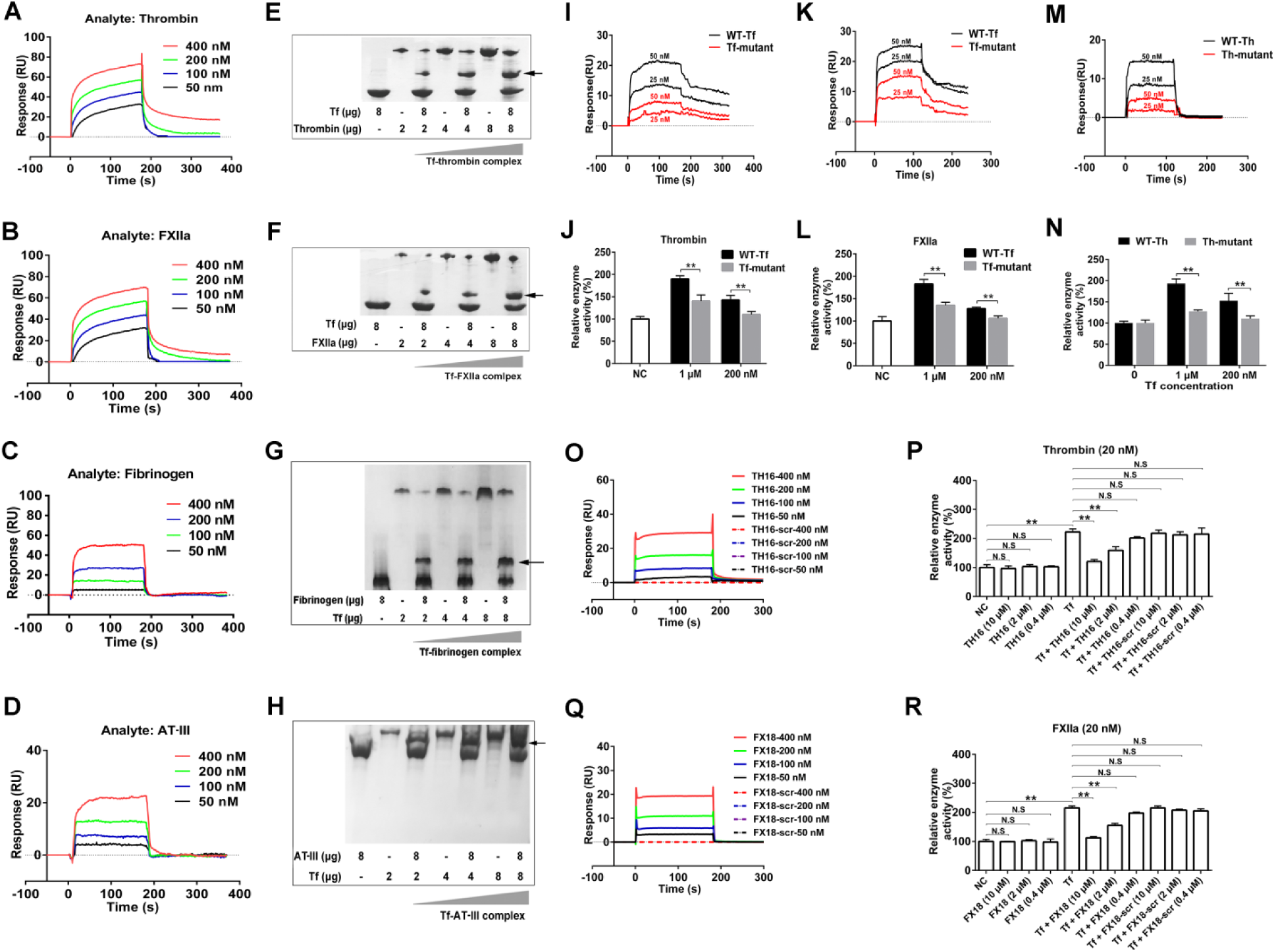
Interactions between Tf and clotting factors. SPR analysis of the interaction between Tf and thrombin **(A)**, FXIIa **(B)**, fibrinogen **(C)** or antithrombin III (AT-III) **(D)**. Native gel shift analysis of the interaction between Tf (8 μg) and thrombin (2, 4 and 8 μg) **(E)** or FXIIa (2, 4 and 8 μg) **(F)**. Native gel shift analysis of the interaction between Tf (2, 4 and 8 μg) and fibrinogen (8 μg) **(G)** or AT-III (8 μg) **(H)**. The arrow indicates the complex of Tf-thrombin, Tf-FXIIa, Tf-fibrinogen or Tf-AT-III. SPR analysis of the interaction between wild-type Tf (WT-Tf) or Tf mutant (E333R and E338R) and thrombin **(I)** or FXIIa **(K)**. The effects of wild-type Tf and Tf mutant on enzymatic activity of thrombin **(J)** and FXIIa **(L)**. SPR analysis of the interaction between Tf and wild-type thrombin (WT-Th) or thrombin mutant (Th-mutant, R117A and R122A) **(M).** The effects of Tf on enzymatic activity of wild-type thrombin and thrombin mutant **(N)**. SPR analysis of the interaction between Tf and TH16 or TH16-scr (scrambled control of TH16) **(O)**, and FX18 or FX18-scr (scrambled control of FX18) **(Q). (P)** Effects of TH16 and TH16-scr on the potentiating activity of Tf on thrombin. **(R)** Effects of FX18 and FX18-scr on the potentiating activity of Tf on FXIIa. Data represent mean ± SD of six independent experiments, \*\**p* < 0.01 by unpaired t-test **(J, L, N)**. \*\**p* < 0.01 by one-way ANOVA with Dunnett’s post hoc test **(P, R)**.

We next determined the actual protein-protein binding properties behind the complexes formation between Tf and thrombin/FXIIa. Docking the modeled structure of Tf-thrombin complex indicated that the conformation of thrombin exosite I is affected by the binding of Tf through several electrostatic interactions and one pool hydrophobic effect (Figure S7). The binding of Tf makes the groove-like exosite I site more open and wide. This conformation change results in a little shift of catalytic triad, which is located adjacent to exosite I. Therefore, substrates of thrombin have more spaces to access the triad residues. Similar to the Tf-thrombin complex, FXIIa interacts with Tf through its own groove-like domain (exosite I domain analogous), which is almost the same folding mode, and changes its conformation as the exosite I of thrombin does (Figure S7).

Based on the docking model and the structural characteristics of the Tf-thrombin/FXIIa complex, we found that two key residues of Tf (E333 and E338) may play key roles in the interaction of Tf with thrombin/FXIIa. The mutant of Tf (E333R and E338R) was thus constructed (Figures S8A-S8E). Notably, the Tf mutant exhibited weak interaction with thrombin (Figure 3I) or FXIIa (Figure 3K). The ability of Tf mutant to potentiate thrombin (Figure 3J) or FXIIa (Figure 3L) was significantly decreased by comparison with wild-type Tf. Furthermore, the docking model also suggested two key residues of thrombin (R117 and R122) play a key role in the interaction of thrombin with Tf. The corresponding mutant (R117A and R122A) of thrombin (Figures S8F-S8H) exhibited significant weak interaction with Tf compared with wild-type thrombin (Figure 3M). Although the enzymatic activity of the thrombin mutant was not influenced, the ability of Tf to potentiate the thrombin mutant was significantly decreased (Figure 3N).

Given that Tf potentiates thrombin and FXIIa by interacting with them, we hypothesize that specific peptides at the binding site of thrombin or FXIIa may competitively bind to the same target in Tf and thus inhibit the effects of it on thrombin and FXIIa. Exosite I motif-based inhibitor peptides TH16 (RIGKHSRTRYERNIEK) and FX18 (RRNHSCEPCQTLAVRSYR) were designed. SPR analysis revealed the critical interaction for the binding specificity of Tf to TH16 (Figure 3O) and FX18 (Figure 3Q). *Ka, Kd* and *KD* for Tf-TH16 was 3.1 × 10^3^ M^−1^s^−1^, 0.3 × 10^−3^ s^−1^ and 97 nM, and that for Tf-FX18 was 2.8 × 10^3^ M^−1^s^−1^, 0.4 × 10^−3^s^−1^ and 143 nM, respectively. However, the scrambled peptide of TH16 (TH16-scr, RKKGIRRYTERHSNIE) and FX18 (FX18-scr, SCPTHYSRQRCRNAVLER) showed no interaction with Tf. Functional study showed that both TH16 and FX18 inhibited the potentiating activity of Tf on thrombin (Figure 3P) and FXIIa (Figure 3R) in a dose-dependent manner. TH16 or FX18 alone showed no effect on the enzymatic activity of thrombin or FXIIa even the concentration up to 10 μM.

### Elevated Tf-prothrombin/thrombin and Tf-FXII/FXIIa complexes in human atherosclerotic plasma and lesions

To confirm whether Tf forms complex with thrombin/FXIIa in atherosclerotic plasma or lesions, we used Bis (sulfosuccinimidyl) suberate (BS^3^) to stabilize the possible components that complex with Tf. Western blot analysis using anti-Tf antibody showed 3 bands (Figure 4A, the top panel) including Tf, and Tf-prothrombin/FXII complexs confirmed by antibodies against prothrombin and FXII (Figure 4A), respectively. The amount of Tf-prothrombin/FXII complexes in CHD patients’ plasma was higher than that in normal controls (Figures 4B and 4C). Co-immunoprecipitation analysis further revealed the formation of Tf-prothrombin and Tf-FXII complexes in plasma (Figure 4D). The presence of Tf- and thrombin-/FXIIa-positive deposits indicated the formation of Tf-thrombin/FXIIa complexes in atherosclerotic plaque (Figure 4E). Moreover, the formation of Tf-prothrombin and Tf-FXII complexes was also observed in atherosclerotic plaque by western blot analysis after being cross-linked, and the amount of which was greater than that in normal controls (Figures 4F-4H).

**Figure 4.**
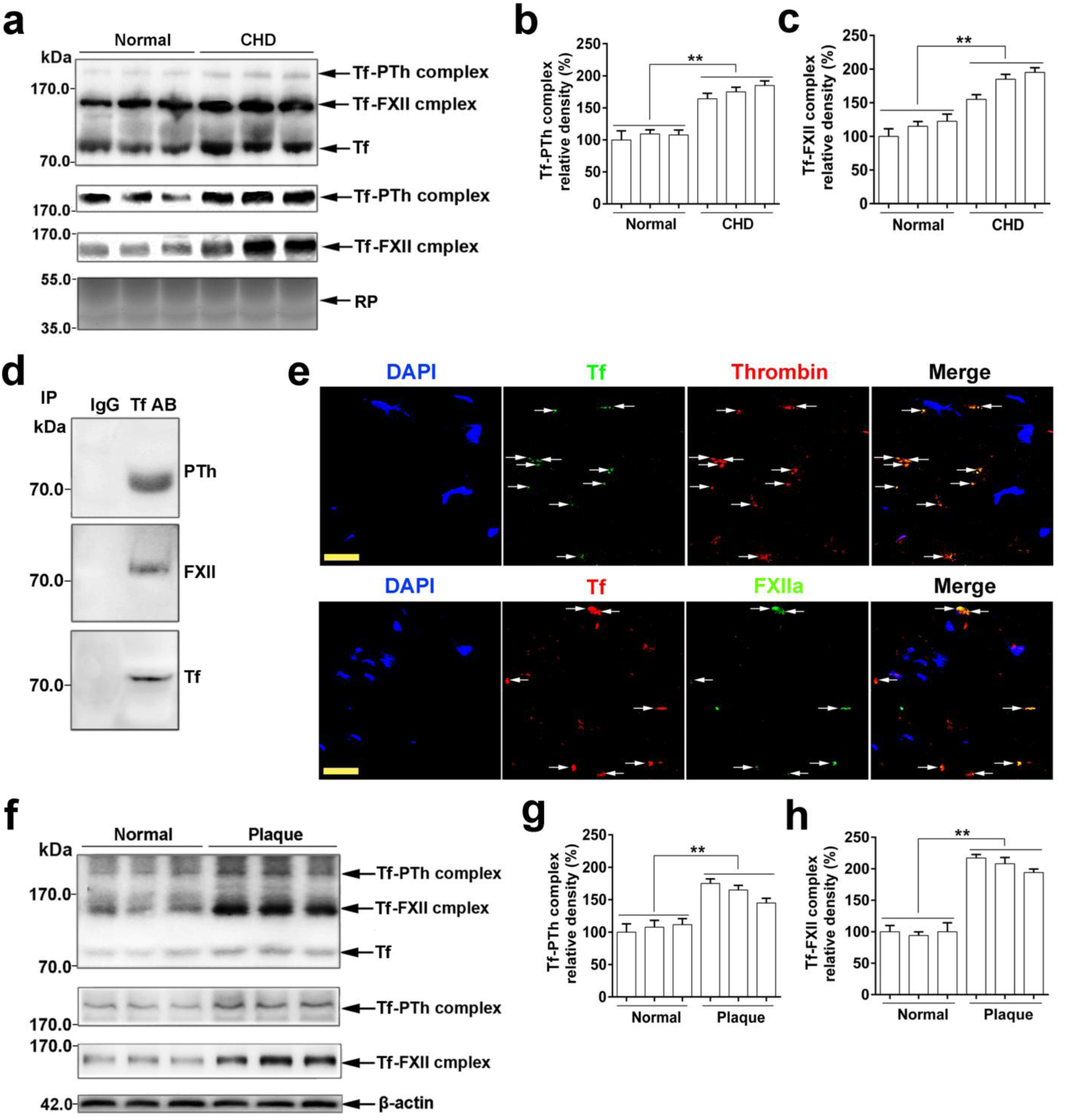
Elevated levels of Tf-thrombin/FXIIa complexes in CHD patient plasma and atherosclerotic plaque. **(A)** Western blot analysis of Tf-prothrombin (Tf-PTh) and Tf-FXII complex in healthy (Normal) and CHD plasma and the quantification of Tf-PTh **(B)** and Tf-FXII **(C)** complex. Red Ponceau (RP)-stained blot was the loading control. **(D)** Co-immunoprecipitation of Tf and prothrombin or FXII in human normal plasma. **(E)** Human atherosclerotic plaque was labeled with either anti-Tf antibody (green) or anti-thrombin antibody (red) to detect the presence of Tf-thrombin complex (top), or labeled with either anti-Tf antibody (red) or anti-FXIIa antibody (green) to detect the presence of Tf-FXIIa complex (bottom). Cell nuclei were labeled by DAPI. Arrows indicate Tf-thrombin- or Tf-FXIIa-positive structures. Scale bar represents 30 μm. Images were representative of at least three independent experiments. Western blot analysis **(F)** and quantification of Tf-prothrombin complex **(G)** and Tf-FXII complex **(H)** in the supernatants of the homogenized thoracic aorta tissue from normal controls and atherosclerotic patients. Data represent mean ± SD (n = 12), \*\**p* < 0.01 by unpaired t-test.

### Tf over-expression aggravates atherosclerotic lesion and hypercoagulability which are attenuated by Tf down-regulation

To elucidate the role of Tf in atherosclerotic lesion development, the effects of Tf over-expression and knockdown on the development of AS and hypercoagulability were evaluated. The Tf expression levels were first validated by qRT-PCR and western blot (Figure S9). After the viruses were injected, the *Apoe*^−*/*−^ mice were fed with a HFD for 6 weeks to spark and study the development of AS and hypercoagulability. As illustrated in Figure 5A, an increase in plasma concentration of Tf was observed in Tf over-expressed *Apoe*^−*/*−^ mice (PLP-Tf) compared with *Apoe*^−*/*−^ controls (NC) or blank virus (with empty over-expression (PLP) vector) injected mice. Conversely, a reduction in plasma concentration of Tf was found in Tf knockdown *Apoe*^−*/*−^ mice (RNR-Tf) in comparison with *Apoe*^−*/*−^ controls or blank virus (with empty knockdown (RNR) vector) injected mice. The plasma from PLP-Tf mice showed significantly elevated activities of thrombin (Figure 5B) and FXIIa (Figure 5C) as analyzed by using synthetic substrates and shortened activated partial thromboplastin time (APTT) (Figure 5D), prothrombin time (PT) (Figure 5E), and tail bleeding time (Figure 5F). In contrast, the plasma from RNR-Tf mice showed significantly reduced activities of thrombin and FXIIa but prolonged APTT, PT, and tail bleeding time (Figures 5B-5F), suggesting that the plasma level of Tf regulates enzymatic activities of thrombin and FXIIa as well as APTT, PT, and tail-bleeding time. No significant changes at iron metabolism indexes and erythrocyte indexes were observed after Tf over-expression or knock-down (Table S3).

**Figure 5.**
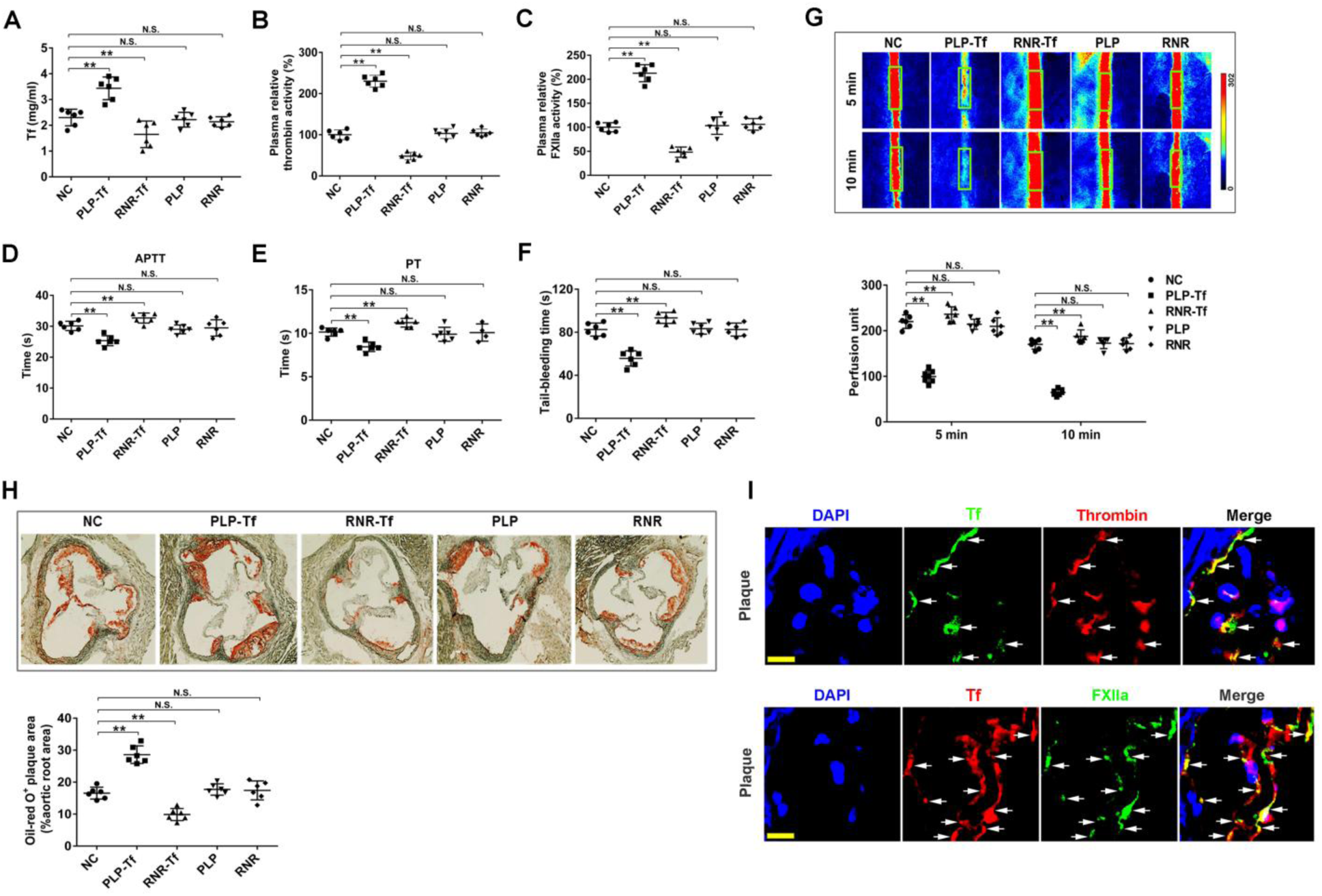
The effects of Tf over-expression and knockdown on atherosclerotic development and hypercoagulability. **(A)** The plasma concentrations of Tf in five groups of *Apoe*^−*/*−^ mice fed by HFD for 6 weeks (Tf over-expression (PLP-Tf) and its blank PLP, knockdown (RNR-Tf) and its blank RNR, and normal *Apoe*^−*/*−^ mice (NC)). Relative activity of thrombin **(B)** and FXIIa **(C)**, APTT **(D)**, PT **(E)** in their plasma and tail-bleeding time **(F)** were also shown. **(G)** Representative images of carotid artery blood flow (top) in FeCl_3_-treated mice by laser speckle perfusion imaging, and the region of interest (the rectangle in green) was placed in the carotid artery to quantify blood flow change. Relative blood flow in the region of interest was shown (bottom). **(H)** The representative images of oil-red O-stained atherosclerotic plaques (top) and quantitative analysis of stained area (bottom) were shown. Data represent mean ± SD (n = 6), \*\**p* < 0.01 by one-way ANOVA with Dunnett’s post hoc test. **(I)** Atherosclerotic plaques from the mice fed by HFD for 4 weeks were labeled with either anti-Tf, anti-thrombin, or anti-FXIIa antibody. Cell nuclei were labeled by DAPI. Arrows indicate Tf-thrombin- or Tf-FXIIa-positive structures. Scale bar represents 30 μm. Images were representative of at least three independent experiments. N.S.: no significance.

The role of Tf in promoting hypercoagulability was further investigated in mouse thrombosis model induced by FeCl_3_. As illustrated in Figure 5G, blood flow in the carotid artery was significantly decreased in Tf over-expressed mice, suggesting an increased thrombus formation. Inversely, blood flow was increased in the Tf knockdown mice, suggesting an inhibited thrombosis formation. Consequently, as illustrated in Figure 5H, a substantial increase in plaque size in the aortic root was observed in Tf over-expressed *Apoe*^−*/*−^ mice, while reduced lesion formation was found in Tf knockdown *Apoe*^−*/*−^ mice. In addition, the Tf- and thrombin/ FXIIa-positive deposits (Figure 5I) were also observed in the atherosclerotic lesion of *Apoe*^−*/*−^ mice. These findings demonstrate that Tf plays a key role in hypercoagulability and atherosclerotic lesion development.

### Interference with Tf exerts anti-hypercoagulability and anti-AS effects

After a 6-week treatment with an anti-Tf antibody, a decrease in plasma concentration of Tf was observed (Figure S10A), which is accompanied with the relative decreased enzymatic activities of thrombin (Figure S10B) and FXIIa (Figure S10C) as well as prolonged APTT (Figure S10D), PT (Figure S10E) and tail-bleeding time (Figure S10F), indicating decreased hypercoagulability in *Apoe*^−*/*−^ mice depleted of Tf. Consequently, an obvious decrease in plaque size was observed in anti-Tf antibody treated *Apoe*^−*/*−^ mice (Figure 6A). Importantly, no statistically significant changes at iron metabolism indexes and erythrocyte indexes were observed after anti-Tf antibody treatment (Table S3).

**Figure 6.**
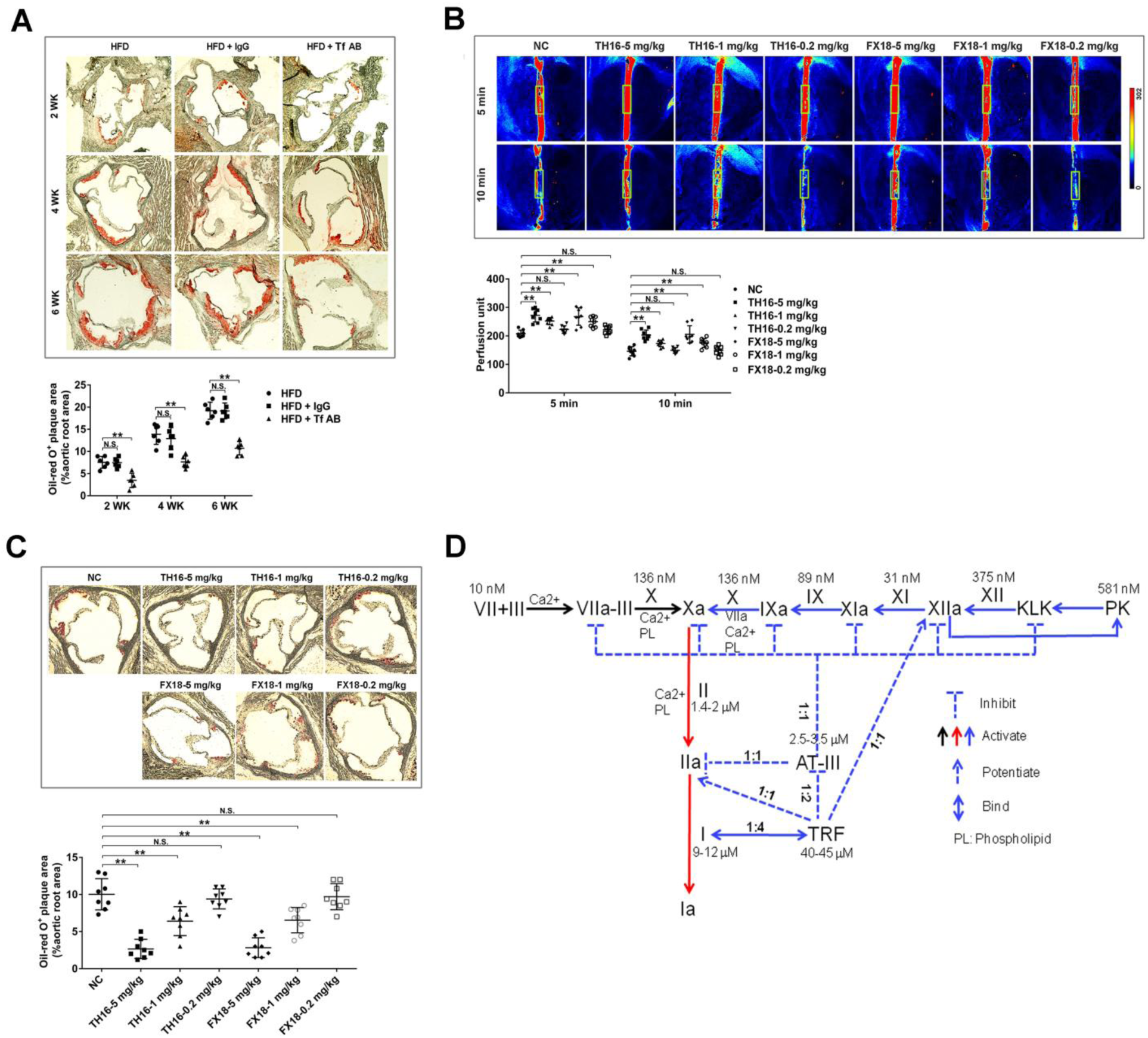
Tf interferences exert anti-AS effects *in vivo*. The HFD-fed *Apoe*^−*/*−^ mice were subjected to anti-Tf antibody (Tf AB) or control IgG treatment 2 times/week for 6 weeks. **(A)** Representative images (top) of oil-red O-stained plaques and quantitative analysis (bottom) of the stained area were shown. **(B)** Effects of TH16 and FX18 on FeCl_3_-induced carotid artery thrombus formation in C57BL/6J mice. Representative images of carotid artery blood flow (top) and quantitation (bottom) were shown. **(C)** Effects of TH16 and FX18 on mouse AS development. Representative images (top) of oil-red O-stained plaques and quantitative analysis (bottom) of the stained area were shown. **(D)** The graphical representation of Tf’s central role and its interactions with clotting factors to maintain coagulation balance. Tf takes part in three types of interactions in the coagulation balance including: 1) most of Tf (TRF, ∼ 40 μM) is sequestered by binding with fibrinogen (∼ 10 μM) with a molar rate of 4:1; 2) Tf blocks inactivation of AT-III towards thrombin and FXa by binding with AT-III with a molar rate of 2:1;3) Tf interacts and potentiates thrombin and FXIIa with a molar rate of 1:1. Data represent mean± SD (n = 6-8), \*\**p* < 0.01 by one-way ANOVA with Dunnett’s post hoc test. N.S.: no significance.

As illustrated in Figures S10G and S10H, both of exosite I motif-based peptides TH16 and FX18, which only inhibited Tf’s potentiating ability on thrombin and FXIIa without direct effect on the enzymes (Figures 3P and 3R), prolonged plasma recalcification time in a dose-dependent manner, suggesting that the coagulation was inhibited by the two peptides. The increase in plasma recalcification time may further contribute to an extended bleeding time in mice (Figure S10I). The inhibitory effects of TH16 and FX18 on thrombosis formation were further demonstrated by an increased blood flow in the carotid artery *in viv*o (Figure 6B). Moreover, after treatment for 3 weeks, TH16 and FX18 significantly decreased atherosclerotic plaque size in a dose-dependent manner, indicating their anti-AS effects (Figure 6C). Together, these data indicate that Tf drives development of atherosclerotic plaque and that interference with Tf might harbor valuable opportunities for AS prevention and treatment.

## Discussion

A balance between clotting and bleeding is always maintained by complicated interactions between coagulation and anti-coagulation system in the body under normal physiology. The coagulation process is under the inhibitory control of several inhibitors that limit the clot formation, thus avoiding the thrombus propagation (Esmon, 2000; Palta et al., 2014). Main natural inhibitor of coagulation enzymes in plasma is AT-III with a concentration of ∼3 μM. Compared with AT-III, coagulation proteases’ concentration (pM to nM) is much lower, suggesting that coagulation proteases’ functions are completely blocked by AT-III in normal circulation. Pro-coagulant state is found in CHD plasma, i.e., AS, suggesting that coagulation balance was interrupted and AT-III’s inactivation on coagulation proteases is inhibited, but the process is unknown. By using AS plasma in this study, we found enhanced enzymatic activity of thrombin and FXIIa in atherosclerotic plasma although total levels of prothrombin and FXII are not different from normal plasma (prothrombin ∼2 μM, FXII ∼0.375 μM) (Mari et al., 2008), and thus the concentration of thrombin and FXIIa should be less than 2 and 0.375 μM, respectively, which is less than AT-III’s concentration of ∼3 μM. Considering that AT-III binds and inactivates coagulation enzymes in a ratio of 1:1, we inferred that another plasma factor may block AT-III to contribute enhanced enzymatic activity of thrombin and FXIIa and to induce pro-coagulant state in AS plasma.

Indeed, we found elevated level of Tf in human atherosclerotic plasma, which is associated with enhanced enzymatic activity of thrombin and FXIIa and pro-coagulant state (Figures 1A-1D and Figures S1-S2). The concentration of Tf in CHD patients’ plasma samples was 46% higher than that in controls (4.173 vs 2.865 mg/ml). Additionally, elevated Tf level was also observed in the plasma and atherosclerotic plaque of AS mice (Figures 1F-1H). Further study indicated that Tf directly interacts with fibrinogen, thrombin, FXIIa and AT-III with different affinity to maintain coagulation balance. Tf’s *KD* for thrombin, FXIIa, fibrinogen and AT-III is 7.7, 13.9, 29 and 524 nM, respectively (Figures 3A-3D). Tf showed ability to potentiate enzymatic activities of thrombin and FXIIa, the two key coagulation proteases. Tf exists in plasma in either holo-Tf or apo-Tf, and about 30% to 40% is present as holo-Tf (Clark, 2008; Haber et al., 2008; Stack et al., 2014). We found both holo- and apo-Tf showed a similar ability to bind thrombin, FXIIa, fibrinogen and AT-III. They also showed similar activity to potentiate thrombin and FXIIa and to block AT-III’s inactivation on thrombin and FXa (Figures 2). The formation of Tf-prothrombin/thrombin or Tf-FXII/FXIIa complexes both *in vivo* and *in vitro* was further confirmed by the analysis of SPR, native-PAGE, immunoprecipitation and laser confocal microscopy (Figures 3A-3H and Figure 4). Importantly, Tf binds with fibrinogen with a molar ratio of 4:1, which is consistent with their ratio of molar concentration in normal plasma (40 and 10 μM) (Figure S5E). In addition, Tf also binds with AT-III, thrombin and FXIIa with a molar ratio of 2:1, 1:1 and 1:1, respectively. It suggests that most of Tf is immobilized by fibrinogen in normal plasma, while up-regulated Tf in AS plasma interacts with and potentiates thrombin and FXIIa and blocks the inactivation of AT-III on thrombin and FXa to induce hypercoagulability (Figure 6D).

Tf’s role as a procoagulant was proven by high thrombotic and atherosclerotic risks in Tf over-expressed mice and alleviated atherosclerotic lesion formation and coagulability in the mice interfered with Tf (Tf knockdown, neutralization with anti-Tf antibody) (Figure 5, Figure 6A and Figures S10A-S10F). The docking model and the structural characteristics of the Tf-thrombin or Tf-FXIIa complex suggest that the binding of Tf to exosite I on thrombin or exosite I domain analogous on FXIIa may affect the steric hindrance of the active site cleft. Based on exosite I domains of thrombin and FXIIa, the inhibitor peptides (TH16 and FX18) were developed. These two peptides showed abilities to inhibit the potentiating activity of Tf on thrombin and FXIIa *in vitro* (Figures 3P and 3R), and alleviated hypercoagulability and atherosclerotic lesion *in vivo* (Figures S10G-S10I, Figures 6B and 6C), further confirming that the site in thrombin or FXIIa for interacting with Tf is located in the domain of exosite I, and exosite I motif-based interference of Tf-thrombin/FXIIa interaction represents a novel strategy for the development of clinical anti-AS medicine.

Collectively, our data have demonstrated that Tf plays a central role to maintain coagulation balance by interacting with coagulation and anti-coagulation factors. The old protein Tf is proven as a clotting factor that we would like name Factor VI. Normally, coagulation balance might be achieved by the two interactions: 1) Tf-fibrinogen with a molar ratio of 4:1, and 2) AT-III-coagulation proteases, i.e., thrombin and FXIIa. Coagulation imbalance is probably caused by abnormally elevated Tf, which blocks AT-III’s inactivation on coagulation proteases by binding to AT-III and interacts with and potentiates thrombin/FXIIa besides its normal interaction with fibrinogen, and thus induces hypercoagulability. Furthermore, factors up-regulating Tf may cause hypercoagulability and Tf interference thus represents a promising therapeutic strategy for it, which will be demonstrated in the accompanying paper.

## Methods

### Specimens of human atherosclerotic plaque and plasma

The Institutional Review Board of Kunming Institute of Zoology (KIZ) and the Yan’an Affiliated Hospital of Kunming Medical University approved this study (KIZ-YA-20150109). All human specimens were collected with the informed consent of the patients prior to the study. Human plasma of patients with CHD (n = 42) and healthy controls (n = 42) were collected from Yan’an Affiliated Hospital of Kunming Medical University (Table S1). In total, 42 subjects with CHD who had angiographically-visible luminal narrowing were selected in the present study. The patients with clinical features of angina pectoris were further diagnosed by coronary angiography. The 42 patients were matched with 42 healthy volunteers as a normal control group with angiographically normal coronary arteries, and no history of hypertension, diabetes mellitus, or hypercholesterolemia disease. Immediately following the blood drawing (1.5 % EDTA-Na_2_ was utilized as an anticoagulant agent), plasma was obtained by centrifuging at 3000 rpm/min for 20 min at 4°C, and stored at −80°C after being sub-packed.

Atherosclerotic plaque specimens were obtained from coronary endarterectomy, and normal arteries were obtained from coronary artery bypass surgery (n = 12, age 30-80) (Table S2) from Yan’an Affiliated Hospital of Kunming Medical University. Immediately following surgical removal, some specimens were minced and homogenized at 4°C for protein extraction, and further stored at −80°C after being sub-packed; some were fixed in 10% buffered formalin to prepare a frozen section; others were placed in RNAlater (R0901-500ML, Sigma, USA) for RNA extraction and stored at −80°C for further use.

### Animals and ethics statement

All animal experiments were approved by the Animal Care and Use Committee at Kunming Institute of Zoology (SMKX-2016013) and conformed to the US National Institutes of Health’s Guide for the Care and Use of Laboratory Animals (The National Academies Press, 8th Edition, 2011). *Apoe*^−*/*−^ mice (female, 8 weeks old, C57BL/6J background, number of backcrosses: 10 times) and C57BL/6J mice (female, 8 weeks old) were purchased from Vitalriver Experiment Animal Company (Beijing, China) and housed in a pathogen-free environment.

### Chromogenic assays

Effects of plasma from CHD patients and healthy controls on proteases (kallikrein, FXIIa, FXIa, FVIIa and thrombin) involved in coagulation were tested using corresponding chromogenic substrates. The testing enzyme was incubated with the plasma (1 μl) in 60 μl Tris-HCl buffer (50 mM, pH 7.4) for 5 min, and a certain concentration of chromogenic substrate was then added as described below. Absorbance at 405 nm was monitored immediately, and the kinetic curve was recorded using an enzyme-labeled instrument (Epoch, BioTek, USA) for 30 min. Relative enzyme activity was obtained by calculating the velocity of enzymatic hydrolysis of its substrate. Human α-thrombin (20 nM, T6884, Sigma, USA) and human α-FXIIa (20 nM, HFXIIa 1212a, Enzyme Research Laboratories, USA) were reacted with 0.2 mM chromogenic substrate of *H-D*-Phe-Pip-Arg-*p*Na·2HCl (CS-01, Hyphen Biomed, France) and *H-D*-Pro-Phe-Arg-*p*NA·2HCl (CS-31, Hyphen Biomed, France), respectively. The used concentration for kallikrein (HPKa 1303, Enzyme Research Laboratory, USA) and FXIa (HFXIa 1111a, Enzyme Research Laboratory, USA) was 400 nM, and the corresponding chromogenic substrate was 0.2 mM *H-D*-Pro-Phe-Arg-*p*NA·2HCl (CS-31, Hyphen Biomed, France). The used concentration for FVIIa (HFVIIa, Enzyme Research Laboratory, USA) was 20 nM, and the chromogenic substrate was 0.1 mM CH_3_SO_2_-D-CHA-But-Arg-*p*NA·AcOH (ADG217L, Sekisui Diagnostics, Germany).

### Purification and identification of Tf

Albumin and IgG were first removed using HiTrap Albumin and IgG depletion column (GE, USA). Plasma was diluted (1:1) by 20 mM Tris-HCl buffer containing 20 mM NaCl (pH 7.8) and then applied to a Resource Q column (17-1177-01, GE, USA) for purification in a fast protein liquid chromatography (FPLC) system (GE, USA). The column was pre-equilibrated with solvent A (20 mM Tris-HCl, pH 7.8), and the elution was performed with a linear gradient of 0-100% solvent B (20 mM Tris-HCl, 1 M NaCl, pH 7.8) over 100 min. The fraction containing potentiating ability on thrombin and FXIIa was subjected to a Mono Q column (17-5166-01, GE, USA) using the same elution system as above for further purification.

The purified thrombin/FXIIa potentiator (10 μg) was dissolved in 25 mM NH_4_HCO_3_ buffer and reduced by 10 mM dithiothreitol (DTT) for 1 h at 37°C. The reduced sample was alkylated by 30 mM iodoacetamide dissolved in the same buffer for 30 min at room temperature in a dark place. The alkylation reaction was ended by adding additional DTT. The sample was treated with 1% trypsin (w/w) at 37°C overnight for mass spectrometry analysis. The matrix-assisted laser desorption ionization time-of-flight mass spectrometer (MALDI-TOF/TOF mass spectrometer, Autoflex speed, BrukerDaltonics, Germany) was employed for data acquisition according to the manufacturer’s instruction. The MS and MS/MS spectra were collected and processed using the FlexControl, FlexAnalysis and BioTools software (BrukerDaltonics, Germany).

### Tf measurement in human plasma and tissue specimens

The concentration of Tf in the plasma of CHD patients and healthy controls was determined using human Tf ELISA kit (EK12012-96T, Multi Sciences, China) according to the manufacturer’s instruction. Amounts of Tf in plasma and plaque homogenates were also determined by western blot analysis. Briefly, plasma and tissue homogenates were first separated by 12% sodium dodecyl sulfate-polyacrylamide gel electrophoresis (SDS-PAGE), and then transferred to polyvinylidene difluoride (PVDF) membranes. An anti-Tf antibody (1:2000, 11019-RP02, Sino Biological Inc, China) was used in the immunoreactivity. To control for plasma loading and transfer, membranes were stained by Red Ponceau after transferring, or blotting for β-actin as a plaque homogenates loading control.

### Effects of Tf on enzymatic activity of coagulation factors

Effects of apo-Tf or holo-Tf (Sigma, purity greater than 98%, no residual enzyme activity of FXIIa and thrombin) on proteases involved in coagulation (kallikrein, FXIIa, FXIa, FVIIa and thrombin) were tested using corresponding chromogenic substrates as described above. Human serum albumin (HSA, purity greater than 98%) was used as a control. The effect of Tf on thrombin to hydrolyze its natural substrate (fibrinogen) was analyzed by a reverse phase high performance liquid chromatography (RP-HPLC) system. Briefly, human α-thrombin (0.1 NIH unit) in 40 μl Tris-HCl (25 mM, pH 7.4) was incubated with 500 μl fibrinogen (10 mg/ml, 16088, Cayman, USA) in same buffer containing 0.15 M NaCl in the presence of apo-Tf (0.2-5 μM) for 30 min at 37°C. After the incubation, 500 μl of 20% trichloracetic acid (TCA) was added to stop the reaction and then centrifuged at 12,000 rpm for 10 min to precipitate insoluble protein. The aliquots of 700 μl of supernatant were used for RP-HPLC analysis. The elution system consisted of solvent A (0.025 M ammonium acetate, pH 6.0) and solvent B (50% acetonitrile in 0.05 M ammonium acetate, pH 6.0) using a linear gradient of 0-100% solvent B over 100 min. Release of FbpA and FbpB was quantified by calculating the corresponding eluted peak area on a C_18_ column (30 cm × 0.46 cm, Hypersil BDS, USA), respectively.

The effect of Tf on FXIIa to hydrolyze its natural substrate (prekallikrein, PK) was assayed by SDS-PAGE. PK (10 μg, HPK1302, Enzyme Research Laboratory, USA) was incubated with human α-FXIIa (0.01 NIH units) in 40 μl of Tris-HCl buffer (50 mM, pH 7.4) in the presence of apo-Tf (0.2-5 μM). After 30 min incubation at 37°C, all of the reactions were applied to 12% SDS-PAGE. The productions of kallikrein heavy chain (HC∼52 kDa) and light chain (LC∼36 and 33 kDa) were detected by western blot using anti-plasma PK polyclonal antibody (1:1000, SAPK-IG, Enzyme Research Laboratories, USA). PK, FXIIa heavy chain (FXIIa HC) and Tf were also detected using anti-plasma PK, anti-FXII (1:2000, ab242123, Abcam, USA) and anti-Tf (1:2000, 11019-RP02, Sino Biological Inc, China) antibodies, respectively. The HC of kallikrein was quantified using Image J software. Tf’s effects on prothrombin (HP 1002, Enzyme Research Laboratories, USA) and FXII (HFXII 1212, Enzyme Research Laboratories, USA) activation were assayed using corresponding chromogenic substrates and reaction system as described above.

### Effect of Tf on thrombin/FXa-AT-III complexes formation *in vitro*

Briefly, Tf (2.5-10 μM), AT-III (2 μM, A2221-125UG, Sigma, USA) and thrombin (20 nM) or FXa (20 nM, HFXa 1011, Enzyme Research Laboratory, USA) were incubated simultaneously in 60 μl Tris-HCl buffer (50 mM, pH 7.4) for 5 min, and thrombin or FXa activity were tested using corresponding chromogenic substrates as described above for thrombin and Z-D-Arg-Gly-Arg-pNA·2HCl (S-2765, Aglyco, China) for FXa. Thrombin or FXa only or incubation with AT-III in the same buffer were also analyzed. Equal HSA was used as a control. Thrombin-AT-III complex (TAT) was tested using ELISA kit (ab108907, Abcam, USA) according to the manufacturer’s instruction. FXa-AT-III complex level was measured by sandwich ELISA as described below. Briefly, plates were coated with anti–human FXa antibody (PAB19898, Abnova, USA) and blocked with 2% BSA before incubation with the above reaction solution. After washing, FXa-AT-III complex was detected by incubation with HRP-conjugated anti-human AT-III antibody (1: 200, SAAT-APHRP, Enzyme Research Laboratory, USA). Relative level of TAT and FXa-AT-III complex was calculated.

### Effects of Tf on blood coagulation

Healthy human plasma was collected from the Kunming Blood Center. To test the effect of Tf on plasma recalcification time, 20 μl of plasma was incubated with human apo- or holo-Tf (0.2-5 μM) in 60 μl of HEPES buffer (20 mM HEPES, 150 mM NaCl, pH 7.4) for 10 min at 37 °C, and 60 μl of 25 mM CaCl_2_ preheated at 37 °C was then added. The clotting was monitored at 650 nm and clotting time was calculated by measuring the time to half maximal increase in absorbance.

### Mouse AS model

*Apoe*^−*/*−^ mice (female, 8 weeks old) were fed a high fat diet (HFD, 21% fat, 0.15% cholesterol) for 6 weeks to induce AS, and some other *Apoe*^−*/*−^ mice were fed a normal diet (ND) as controls. After the 6-week induction, the mice were sacrificed and the blood were collected for further plasma preparation (blood was obtained by mixing 1 volume trisodium citrate (0.13 M) with 9 volumes blood, the plasma was then obtained by centrifuging at 3000 rpm/min for 20 min at 4 °C). The aortic root was immediately fixed in 4% paraformaldehyde dissolved in PBS at 4 °C overnight following surgical removal, and organs including liver, brain, spleen, muscle, kidney, stomach, and plaque were also collected and placed in RNAlater for RNA extraction or PBS (20 mM phosphate, pH 7.4, 100 mM NaCl) for protein extraction. The mouse aortic root specimens were cut into 8 μm sections using a freezing microtome (Thermo Scientific). Some sections of aortic root were stained by oil-red O to investigate atherosclerotic lesion, and some others were used for immunofluorescent assay as described below. Images of oil-red O staining were acquired by a dark field microscope (Life technologies), and the oil-red O stained plaque area was measured by Image J.

The RNA extraction and cDNA reverse transcription were performed by using RNA extraction kit (DP419, Tiangen, China) and reverse transcription kit (A5000, promega, USA) according to the manufacturer’s instructions, respectively. Tf expression was quantified by both qRT-PCR (forward primer (5’-3’): GGACGCCATGACTTTGGATG; reverse primer (5’-3’): GCCATGACAGGCACTAGACC) and western blot. PCR was performed on the Bio-Rad CFX-96 Touch Real-Time Detection System. The concentration of Tf in mouse plasma was determined using mouse Tf ELISA kit (ab157724, Abcam, USA). Amount of Tf in plaque homogenates was also determined by western blot analysis using the anti-Tf antibody (1:2000, 11019-RP01, Sino Biological Inc, China) as described above.

### Surface plasmon resonance (SPR) analysis

BIAcore 2000 (GE, USA) was used to analyze the interaction between Tf and clotting factors by using HSA as a control. Briefly, apo-Tf or holo-Tf was first diluted (20 μg/ml) with 200 μl sodium acetate buffer (10 mM, pH 5) and then flowed across the activated surface of the CM5 sensor chip (BR100012, GE, USA) at a flow rate of 5 μl/min, reaching a resonance unit (RU) of ∼2000. The remaining activated sites on the chip were blocked with 75 μl ethanolamine (1 M, pH 8.5). Serial concentrations of thrombin, FXIIa, fibrinogen, AT-III, prothrombin or FXII in Tris-HCl buffer (20 mM, pH 7.4) were applied to analyze the interaction with immobilized apo- or holo-Tf with a flow rate of 10 μl/min. The *KD* (equilibrium dissociation constant) for binding as well as association (*Ka*) and dissociation (*Kd*) rate constants were determined by the BIA evaluation program (GE, USA). CM5 sensor chip was coupled with 100 RU of Tf as the method above. The RU change is reported in BIAcore technology where a 1000 RU response is equivalent to a change in surface concentration of about 1 ng/mm^2^ of protein and the binding molar ratio between Tf and fibrinogen or AT-III was calculated by comparing the saturation RU of flowing phase and RU of immobilized phase as the method described (Liu and Wilson, 2010).

### Native PAGE

Basic native PAGE was used to further analyze the interaction between Tf and clotting factors. In brief, human apo-Tf (8 μg) was first incubated with different concentration of human thrombin or human FXIIa (2-8 μg) in 30 μl Tris-HCl buffer (50 mM, pH 7.4) for 10 min, and then applied to 8% precast gel (PG00810-N, Solarbio, China) to analyze the complex formation between Tf and thrombin/FXIIa in running buffer (0.05 M Trizma, 0.38 M glycine, pH 8.9) at 200 V constant voltage for 1 h. Fibrinogen (8 μg) or AT-III (8 μg) and different concentration of apo-Tf (2-8 μg) were also used to analyze the interaction by basic native PAGE. Staining analysis was performed with the staining solution (0.25% (v/v) Coomassie Brilliant Blue R containing 50% (v/v) methanol, 40% (v/v) dH_2_O and 10% (v/v) acetic acid) and the destaining solution (mainly consists of 15% (v/v) methanol, 10% (v/v) acetic acid and 75% (v/v) dH_2_O).

### Confocal microscopy

Human atherosclerotic plaque or mouse aortic root specimens were first cut into 8 μm sections using a freezing microtome (Thermo Scientific). For immunofluorescence detection to detect Tf, thrombin or FXIIa in the atherosclerotic plaque, the section was first incubated with anti-mouse Tf (1:200, 11019-RP01), anti-human Tf (1: 200, 11019-RP02), anti-thrombin (1:200, ab17199, Abcam, USA), and anti-mouse or human FXII (1:200, SC-6670 or SC-66752, Santa Cruz, USA) antibody at 4 °C overnight, respectively. After washing three times by PBS to remove excess primary antibody, the section was incubated with a fluorescently labeled secondary antibody for 1 h at 37 °C. Cell nuclei were stained with 4, 6-diamidino-2-phenylindole (DAPI, Cat# P36941, Lifetechnologies, USA). Images of immunofluorescence were obtained by an Olympus FluoView 1000 confocal microscope according to manufacture’s instructions.

### APTT and PT assays

For APTT assay, 50 μl of APTT reagent (F008-1, Nanjing Jiancheng Bioengineering Institute, China) was incubated with 50 μl plasma for 3 min at 37°C, and then 50 μl of CaCl_2_ (25 mM) preheated at 37°C was added to test the clotting time by monitoring the absorbance at 405 nm using a semi-automatic coagulation analyzer (ThromboScreen 400c, pacific hemostasis, USA). To test PT, 50 μl plasma preheated at 37°C was mixed with 100 μl PT reagent (F007, Nanjing Jiancheng Bioengineering Institute, China) preheated at 37°C for 15 min by monitoring the absorbance at 405 nm.

### Bleeding time measurement

Mouse tail transection model was used to test the tail-bleeding time. In brief, mice tail was cut 2 mm from the tip, and then carefully immersed in 20 ml of saline warmed at 37°C. The bleeding time was recorded until the stream of blood ceased.

### Molecular docking and inhibitory peptides designing

Tf was docked into the exosite I of human thrombin. The model of human thrombin was homologically constructed from known structures (PDB ID: 4NZQ). With the assistance of another two complex structures with PDB ID 1A2C and 1HAH, the binding site was locally placed for further docking processes. Meanwhile, two structures (PDB ID 4XE4 and 4XDE) were used to build the working model of human FXIIa. This FXIIa model was aligned along the thrombin model to confirm the possible groove-like binding area. The human Tf model was then extracted from complex (PDB ID: 3V8X), and this model was cleaned and reconstructed by repairing those missed residues. All of the above three models were optimized by short molecular dynamics to eliminate steric clashes by side chain packing. The modified Tf model was docked into exosite I site of human thrombin and a similar site of FXIIa by the alignment of these two structures. The docking processes were performed with a standard pipeline protocol of Discovery Studio (version 3.1). During the docking process, common parameters were used to obtain more accurate results and ZRank scored the top 2000 poses with RMSD cutoff 10.0 Angstroms.

Based on molecular docking analysis of Tf and thrombin/FXIIa, two peptides (TH16 and FX18) were characterized and synthesized. TH16 (RIGKHSRTRYERNIEK) and FX18 (RRNHSCEPCQTLAVRSYR) were deduced from the sequence of exosite I domain of thrombin (GenBank number (NM_000506.3)) and the sequence of exosite I domain analogous of FXIIa (GenBank number (NP_000496.2)), respectively. The scrambled peptides of TH16 (TH16-scr, RKKGIRRYTERHSNIE) and FX18 (FX18-scr, SCPTHYSRQRCRNAVLER) were also designed and synthesized.

### Effects of Tf interferences on mouse AS development

Several methods of Tf interference, which include lentivirus or retrovirus (10^7^ transducing units) injection into *Apoe*^−*/*−^ mice from tail vein to induce Tf over-expression or knock-down, Tf antibody intravenous injection for two times (50 μg per time) per week for 6 weeks, and inhibitory peptides (TH16 and FX18) intravenous injection with certain dosage (0.2-5 mg/kg) for three times per week for 3 weeks, were performed to evaluate their effects on the development of mouse AS induced by HFD as described above. All the Tf interferences were started from the beginning of HFD induction. According to the methods above, tail-bleeding time, FeCl_3_-induced carotid artery thrombus formation, plasma thrombin and FXIIa generation, APTT, and PT were also evaluated in these mouse models. The amounts of iron and ferritin in plasma were performed using iron test kit (Ferrozine method, TC1016-100T, leagene, China) and ferritin test kit (SEA518Mu, Uscn, China), respectively. Mean corpusular volume (MCV), mean corpusular hemoglobin (MCH) and mean corpuscular hemoglobin concentration (MCHC) were assayed using a blood routine test machine (BC-2800Vet, Mindray, China).

### FeCl_3_-induced carotid artery thrombus formation

The C57BL/6J mice (female, 8 weeks old) or *Apoe*^−*/*−^ mice (female, 8 weeks old) with Tf interference were first anaesthetized with 2.0% isoflurane and core body temperature was maintained at 37 °C during the whole surgery. One of the carotid arteries was exposed by cervical incision, and separated from the adherent tissue and vagus nerve. Thrombosis was induced by applying a piece (2 × 2 mm) of filter paper that was pre-soaked with 10% (w/v) FeCl_3_ solution to the exposed mice carotid artery. The blood flow of the carotid artery of all groups was measured by laser speckle perfusion imaging (PeriCam PSI, HR, Sweden) at 5 min and 10 min after FeCl_3_ induction, respectively. The perfusion unit of region of interest (ROI) was also recorded to quantify the blood flow changes.

### Statistical analysis

The data obtained from independent experiments are presented as the mean ± SD. All statistical analyses were two-tailed and with 95% confidence intervals (CI). The results were analyzed using one-way ANOVA with Dunnett’s post hoc test or unpaired t-test by Prism 6 (Graphpad Software) and SPSS (SPSS Inc, USA). Differences were considered significant at p < 0.05.

## Acknowledgments

We thank Dr. Lin Zeng for technical advice and assistance on MS/MS spectra analysis. This work was supported by funding from Chinese Academy of Sciences (XDB31000000, KFJ-BRP-008 and QYZDJ-SSW-SMC012), National Science Foundation of China (331372208), and Yunnan Province (2015HA023) to R.L. and from the National Science Foundation of China (31640071, 31770835 and 81770464), Chinese Academy of Sciences (XDA12020334 and Youth Innovation Promotion Association).

## Author contributions

X.T., Z.Z., M.F., Y.H., S.W., M.X. and Q.L. performed the experiments and data analysis; Y.L., L.Z., J.W., and B.Y. collected the plasma of atherosclerotic patients; R.L., and X.D. conceived and supervised the project and prepared the manuscript. All authors contributed to the discussions.

## Supplementary information

Supplementary information includes full methods, Figure S1-S10 and Table S1-S3.

