## Supplemental file for "Transferrin plays a central role to maintain coagulation balance by interacting with clotting factors" (from another version of this preprint: version 1, 2019-05-24)

**Materials and Methods**

**Measurement of prothrombin and FXII concentration**

Proteins in the plasma of coronary heart disease (CHD) patients and healthy controls were first separated by sodium dodecyl sulfate-polyacrylamide gel electrophoresis (SDS-PAGE) with 12% gel concentration, and then transferred to polyvinylidene difluoride (PVDF) membranes. The membrane was blocked with 5% non-fat dried milk (BD, USA) dissolved in TBST buffer (2.42 g/L Tris base, 8 g/L NaCl, 0.1% Tween-20, pH 7.6) for 2 h at room temperature. After washing 3 times by TBST buffer, the PVDF membrane was incubated with the primary antibody at 4°C overnight, and then it was incubated with the secondary antibody for another 1 h at room temperature. After washing with TBST, the membrane was developed with an enhanced chemiluminescence kit (PA112, Tiangen, China) by ImageQuant LAS 4000 mini (GE Healthcare, USA). Two primary antibodies (a mouse polyclonal antibody against thrombin (1:2000, ab17199, Abcam, USA) and a goat polyclonal antibody against FXII (1:2000, SC-66752, Santa Cruz, USA)) were used to detect prothrombin and FXII, respectively. To control for plasma loading and transfer, membranes were stained by Red Ponceau after transferring.

**Effects of Tf on thrombin-induced platelet aggregation**

Platelets from healthy volunteers were collected from Kunming Blood Center, Yunnan Province of China. The platelets were washed 3 times by Tyrode's buffer (12 mM NaHCO_3_, 138 mM NaCl, 5.5 mM glucose, 2.9 mM KCl, 2 mM MgCl_2_, 0.42 mM NaH_2_PO_4_, 10 mM HEPES, pH 7.4), and re-resuspended in the same buffer. Platelets (300 μl, 1 × 10^9^ cells/ml) were incubated with Tf (0.2-5 μM) for 5 min at 37°C, and 1 NIH unit of human α-thrombin (T6884, Sigma, USA) was then added to induce platelet aggregation. Light transmittance was monitored immediately and the kinetic curve was recorded using platelet aggregation instrument for 5 minutes (LBY-NJ4, Precil, China). The maximum platelet aggregation rate (MPAR) was calculated.

**Recombinant expression of Tf, thrombin and their mutants**

The prokaryotic expression vector was constructed by inserting DNA sequence encoding mature Tf (GenBank: AAA61140.1, 679 amino acids) or Tf mutant (E333R and E338R) between KpnI and XhoI sites of pET-32a (+) vector. DNA encoding thrombin (NM_000506.3, 308 amino acids) or thrombin exosite I mutant (R117A and R122A) was inserted between BamHI and XhoI sites of pSmart-I vector. The vectors were trasformed into *Escherichia coli* (*E. coli*) Rosetta (DE3), which was induced by 0.8 mM Isopropyl β-D-Thiogalactoside (IPTG) for 6 h in a 110-rpm shaker at 28°C. After the induction, *E. coli* cells were collected by centrifuging at 12000 rpm/min for 15 min at 4°C, and resuspended in the binding buffer (20 mM Tris-HCl, 500 mM NaCl, 5 mM imidazole, pH 7.4). The cells were then homogenized using the Ultrasonic Cell Disruption System (XINYI-IID, XinYi, China). The supernatant was collected by centrifuging at 12000 rpm/min for 2 h at 4°C. Ni^2+^ affinity chromatography column was equilibrated in advance with the binding buffer. The collected supernatant containing fusion protein was subsequently loaded on the Ni^2+^ affinity chromatography column at a flow rate of 0.7 ml/min. The bound fusion proteins were eluted with 5 column volumes of elution buffer (20 mM Tris-HCl, 250 mM NaCl, 500 mM imidazole, pH 7.4). The eluted fraction was resuspended in 20 mM Tris-HCl, pH 7.8, and the salt was removed using an ultrafiltration device (UFC500324, Millipore, USA).

Enterokinase (9014-74-8, C&M Biolabs, USA) was added to the reaction buffer (10 mM Tris, 10 mM CaCl_2,_ pH 8.0) and kept for 24 h at 37°C to release the recombinant Tf. For the release of recombinant thrombin, small Ubiquitin-Like Modifier (SUMO) protease (1 unit, 12588-018, Lifetechnologies, USA) was added to the reaction buffer (50 mM Tris-HCl, 0.2% Igepal, 1 mM dithiothreitol (DTT), pH 8.0) and kept for 16 h at 4°C. Hydrolyzed recombinant proteins were purified by a Mono Q^TM^ 5/50 GL column (17-5166-01, GE, USA) on a fast protein liquid chromatography (FPLC) system. The column was pre-equilibrated with solvent A (20 mM Tris-HCl, pH 7.8) and the elution was performed with a linear gradient of 0-100% solvent B (20 mM Tris-HCl, 1 M NaCl, pH 7.8) over 100 min.

**Crosslinking assay**

Bis (sulfosuccinimidyl) suberate (BS^3^, 1 mM, 21586, Thermo, USA) was used to crosslink plasma (1 μl, CHD patients and normal controls) in HEPES buffer (20 mM, pH 7.4) for 30 min at room temperature. Tris-HCl (10 mM, pH 7.5) was then used to stop the reaction. The crosslinked plasma samples were subjected to 12% SDS-PAGE separation. A rabbit polyclonal antibody against Tf (1:2000, 11019-RP02, Sino Biological, Inc. China) was used to detect Tf. After stripping the PVDF with the stripping buffer (PH1498, Phygene, China), the anti-thrombin (1:2000, ab17199, Abcam, USA) and anti-FXII (1:2000, SC-66752, Santa Cruz, USA) antibody were used to detect prothrombin and FXII, respectively. To control for plasma loading and transfer, membranes were stained by Red Ponceau after transferring. Typical tissue specimens (AS patients, and normal controls) were also crosslinked by BS^3^ and applied for 12% SDS-PAGE to analyze Tf-prothrombin/FXII complexes.

**Immunoprecipitation**

Anti-Tf antibody (2 μg) and human normal plasma (1 μl) were first mixed and incubated for 16 h at 4°C in 30 μl of Tris-HCl buffer (25 mM, pH 7.4). Protein A agarose (20 μl, P2006, Beyotime, China) was then added to couple the anti-Tf antibody and incubated for 3 h at 4°C. After centrifuging at 2500 rpm for 5 min, loading buffer (10 μl, 4 × CW0027A, CWBIO, China) was added and boiled for 10 min to obtain the coupled proteins. All the proteins were subjected to a 12% SDS-PAGE separation and the polyclonal antibody against thrombin (1:2000, ab17199, Abcam, USA), FXII (1:2000, SC-66752, Santa Cruz, USA) and Tf (1: 2000, 11019-RP02, Sino Biological Inc, China) was used to identify prothrombin, FXII and Tf, respectively. IgG (2 μg, NI01, Sigma, USA) was used as a control.

**Generation of lentiviral and retroviral vectors and virus package for Tf over-expression or knockdown**

The coding region of mouse Tf (GenBank: BC092046.1) was synthesized and cloned into pLVX-Puro lentiviral plasmid (Clontech, USA) between EcoRI and BamHI sites. The empty pLVX-Puro plasmid was used as blank control. The oligonucleotides of shRNA sequence targeting Tf (sh-Tf) and its scrambled control shRNA (sh-scr) sequence were synthesized by Sangon Biotech (Shanghai, China). The sequence of sh-Tf is 5’-AAGAATCTGAAGCAGGAAGAC TTCAAGAGAGTCTTCCTGCTTCAGATTCTTCTTTTTT-3’ (Forward oligo), while that for sh-scr is 5’- ACTGAGACTGAACAAAGGAAGTTCAAGAGACTTCCTTTGTTCAGTCTCAGTCTTTTTT-3’ (Forward oligo). The oligonucleotides of sh-Tf and sh-scr including the digested sites of BamHI and EcoRI were annealed with their complementary sequences and inserted into BamHI and EcoRI site of the RNAi-Ready pSIREN-RetroQ retroviral vector (Clontech, USA) to form retroviral vectors of psh-Tf and psh-scr, respectively. Lentiviral vector for Tf over-expression and retroviral vector for Tf knockdown were transfected into murine embryonic liver cells (BNL CL.2 cells, Conservation Genetics CAS Kunming cell Bank, China) to test the efficiency of Tf over-expression or knockdown using western blot and qRT-PCR. The primers for PCR were: forward primer (5’-3’): GGACGCCATGACTTTGGATG and reverse primer (5’-3’): GCCATGACAGGCACTAGACC). The transfection procedure was performed according to the lipofectamine 2000 (11668019, Life technologies, USA) manufacturer's instructions.

HEK 293T cells (Conservation Genetics CAS Kunming cell Bank, China) and EcoPack™ 2–293 cells (Clontech, USA) were used to package lentiviruses and retroviruses, respectively. For each virus preparation, 1.5 × 10^7^ HEK 293T cells or EcoPack™ 2–293 cells were seeded in a T75 flask for 24 h before transfection. pLVX-Puro empty lentiviral vector (20 μg) or with mouse Tf gene and the packaging plasmids including psPAX2 (15 μg, Addgene Inc., USA) and pMD2G (10 μg, Addgene Inc., USA) were co-transfected into HEK 293T cells by lipofectamine 2000. Each flask of EcoPack™2–293 cells was transfected with 25 μg  retroviral vector psh-Tf or psh-scr in 70 μl lipofectamine 2000. The medium was changed with fresh medium after 8 h transfection, and the supernatants containing viruses were harvested after 48 h from the first transfection and concentrated by ultracentrifugation using a P28S rotor (Himac, Japan) at 25,000 rpm for 3 h at 4°C, and then immediately stored at -80°C after being sub-packed. HEK 293T cells (100 μl, 4 × 10^4^ cells) were seeded in a 96-well plate to measure virus titer. Briefly, after the infection by six gradients of (dilution ratio: 10, 10^2^, 10^3^, 10^4^, 10^5^, and 10^6^) lentiviruses (pLVX-Puro-Tf, and negative control) or retroviruses (psh-Tf, and psh-scr) for 48 h, puromycin (2 μg/ml, ab141841, Abcam, USA) was added to screen infected cells for 48 h and virus titer was measured by infection cells (living cells) counting.

**Anti-Tf polyclonal antibody preparation**

Mouse Tf (T0523, Sigma, USA) was used to immunize rabbit to produce anti-Tf polyclonal antibody. Briefly, rabbits (male, 2 kg) were primed by subcutaneous injections of 500 μg mouse Tf in 1 ml complete Freund’s adjuvant (Sigma-Aldrich Inc.) on day 0, followed by subcutaneous booster injections of the half dose of the antigens in incomplete Freund’s adjuvant (Sigma-Aldrich Inc.) on day 14, 28, and 42. The titer and specificity of polyclonal antibodies were screened by indirect ELISA. Anti-Tf polyclonal antibodies were then purified from rabbit serum on a protein G column from Amersham Biosciences (Piscataway, NJ).

**Measurement of thrombin and FXIIa generation of mice plasma**

Plasma from Tf over-expressed, knockdown or anti-Tf antibody treated mice were used in this assay. Briefly, plasma (80 μl) was added into wells of 96-well microtiter plate, and then 20 μl HEPES buffer (20 mM HEPES, 150 mM NaCl, 60 mg/ml BSA, pH 7.4) containing tissue factor (1 pM) (ab87476, Abcam, USA) and phospholipids (4 μM) (20 mol% phosphatidylethanolamine, 20 mol% phosphatidylserine and 60 mol% phosphatidylcholine, all from Sigma, USA) was added. Twenty microliter of thrombin fluorogenic substrate (0.8 mM, Z-Gly-Gly-Arg-AMC, Bachem, Switzerland) containing 20 mM CaCl_2_ was then added to start the reaction. Fluorescence was measured at 30 s intervals during 30 min using microtiter plate fluorometer (Ascent reader, Helsinki, Finland). The excitation and emission wavelengths were 390 and 460 nm, respectively. Relative thrombin activity was calculated by comparing fluorescence values. For plasma FXIIa generation test, briefly, plasma (80 μl) was mixed with 20 μl HEPES buffer, and then 20 μl FXIIa fluorogenic substrate (0.8 mM, Boc-Gln-Gly-Arg-AMC, Bachem, Switzerland) containing 20 mM CaCl_2_ and 40 μg/ml collagen (C9879, Sigma, USA) was added to start the reaction. Relative FXIIa activity was also calculated by comparing fluorescence values.

**Supplementary Figures and Figure legends**

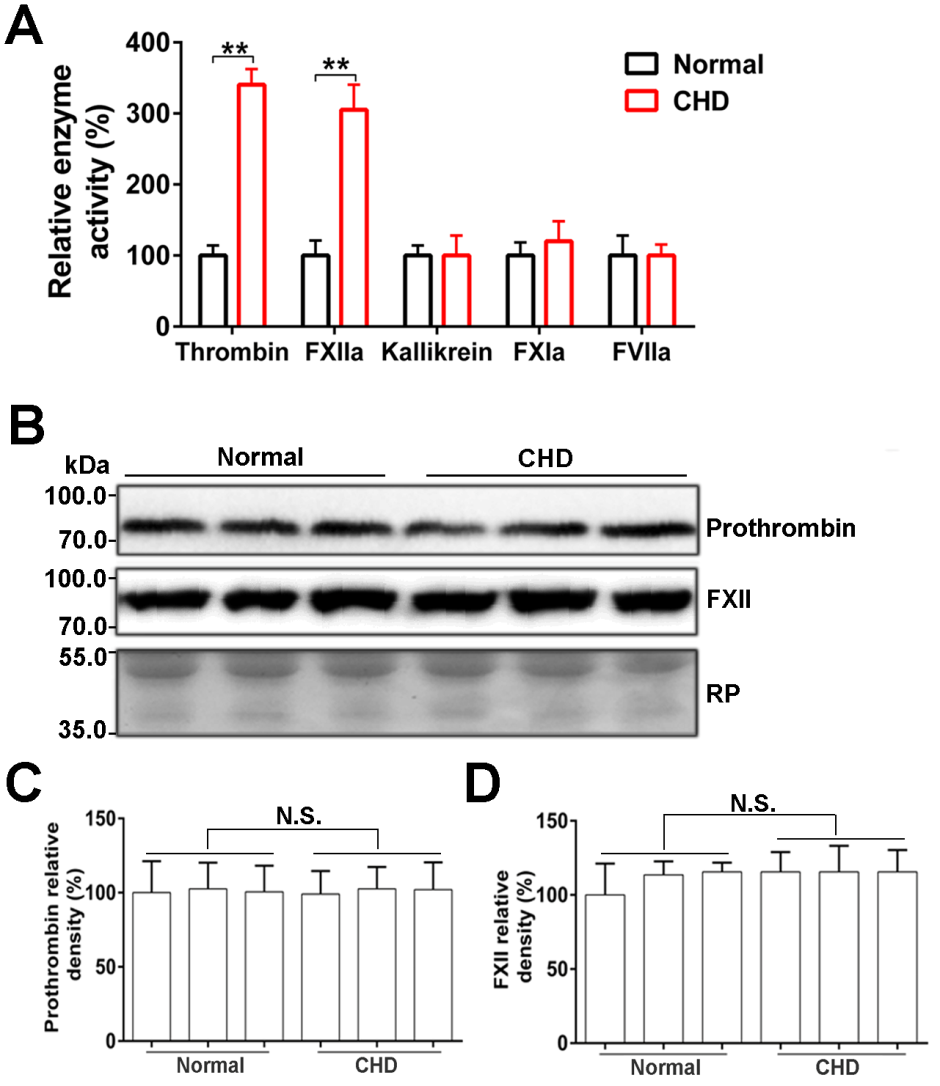

**Figure S1.** **(A)** Effects of plasma from healthy volunteers and CHD patients on enzymatic activities of several coagulants. Western blot **(B)** and quantification analysis of prothrombin **(C)** and FXII **(D)** in plasma of patients. Red Ponceau (RP)-stained blot in panel **b** is the loading control. Data represent mean ± SD (n = 12), ***p* < 0.01 by unpaired t-test. N.S.: no significance.

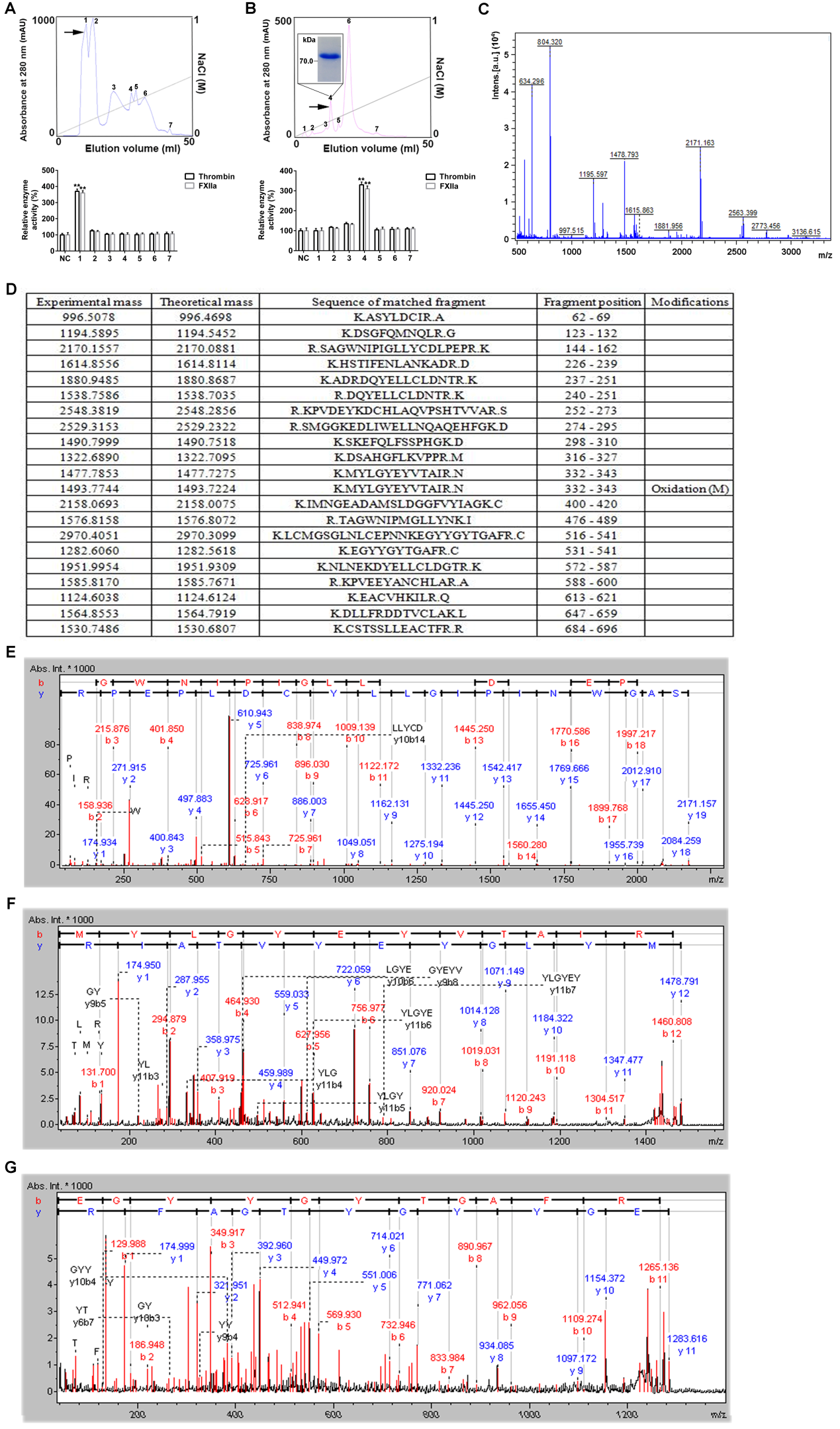

**Figure S2.** **Purification and identification of Tf from CHD plasma.** **(A)** FPLC Resource Q chromatogram of the atherosclerotic plasma after albumin and IgG depletion (top). The main FPLC fractions were applied to test their effects on enzymatic potentiation of thrombin and FXIIa (bottom). **(B)** The pooled fraction indicated by an arrow in panel **(A)** was further purified by a Mono Q FPLC column (top) and pharmacological tests (bottom). Fraction 4 containing the activity to potentiate thrombin and FXIIa was subjected to 12% SDS-PAGE (inset). The thrombin and FXIIa potentiating peaks in **(A)** and **(B)** were indicated by arrows. **(C)** Analysis of trypsin digested products of purified potentiator of thrombin and FXIIa by MALDI-TOF MS. **(D)** The comparison of experimental molecular mass of peptides resulting from trypsin digestion with theoretical molecular mass in NCBI and SwissProt database with Mascot search software. Three peptides produced by trypsin digestion with m/z of 2171.163 **(E),** 1478.793 **(F)** and 1283.613 **(G)**, respectively were selected as parent ions for tandem mass spectrometry. MS/MS spectra of them was used for sequence analysis by BioTools software (Bruker Daltonics, Germany).

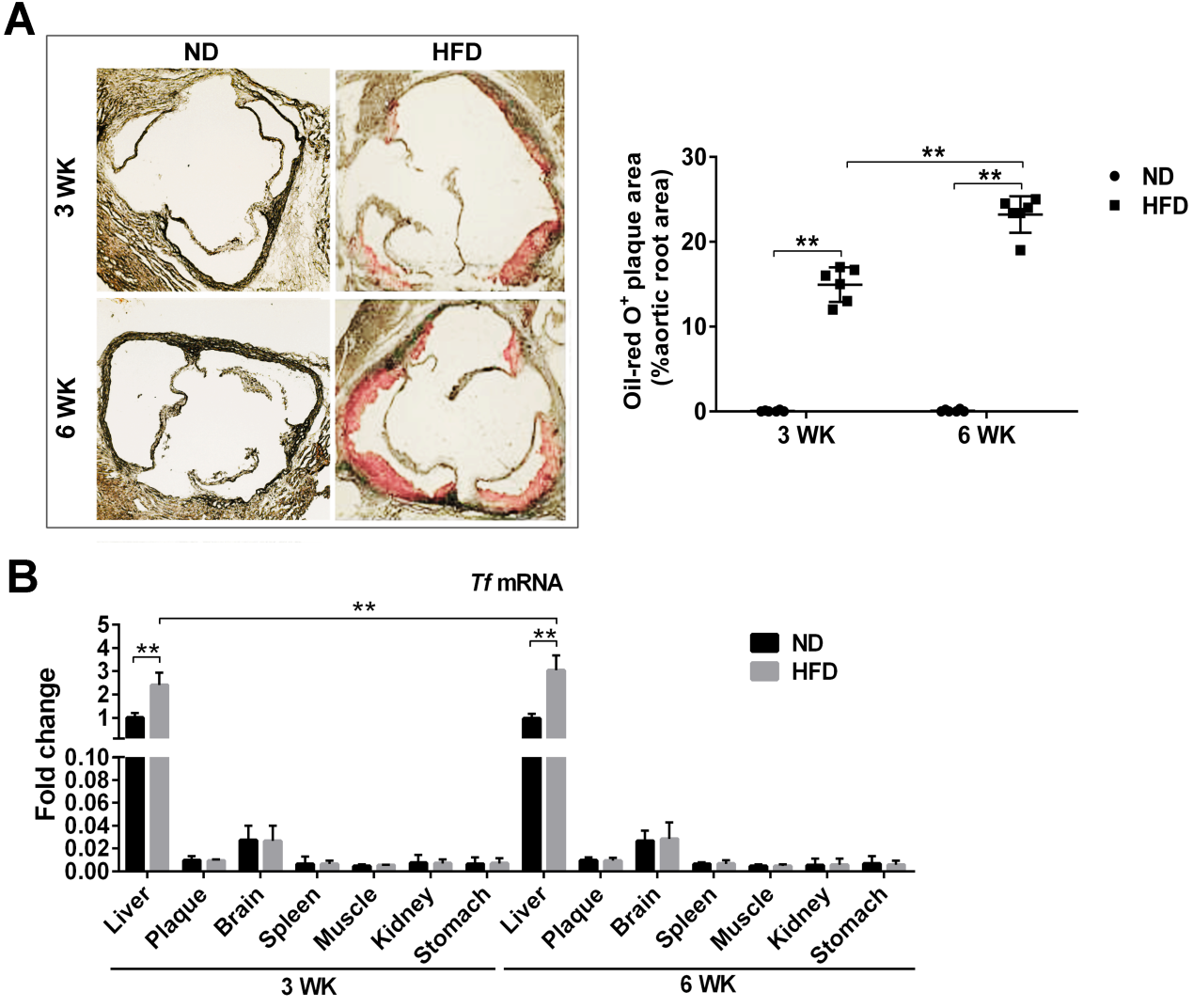

**Figure S3.** **HFD-induced atherosclerosis and tissue distribution of Tf mRNA in mice.** **(A)** *Apoe^-/-^* mice were fed with normal diet (ND) or high fat diet (HFD) for 3 or 6 weeks (WK), and the representative images of oil-red O-stained plaques in arotic root area (left) and quantitative results of stained area (right) were shown. **(B)** Tf mRNA expression in different tissues (liver, plaque, brain, spleen, muscle, kidney, and stomach) was detected by qRT-PCR. Animal experiments were repeated two independent times. Data represent mean ± SD (n = 6), ***p* < 0.01 by unpaired t-test.

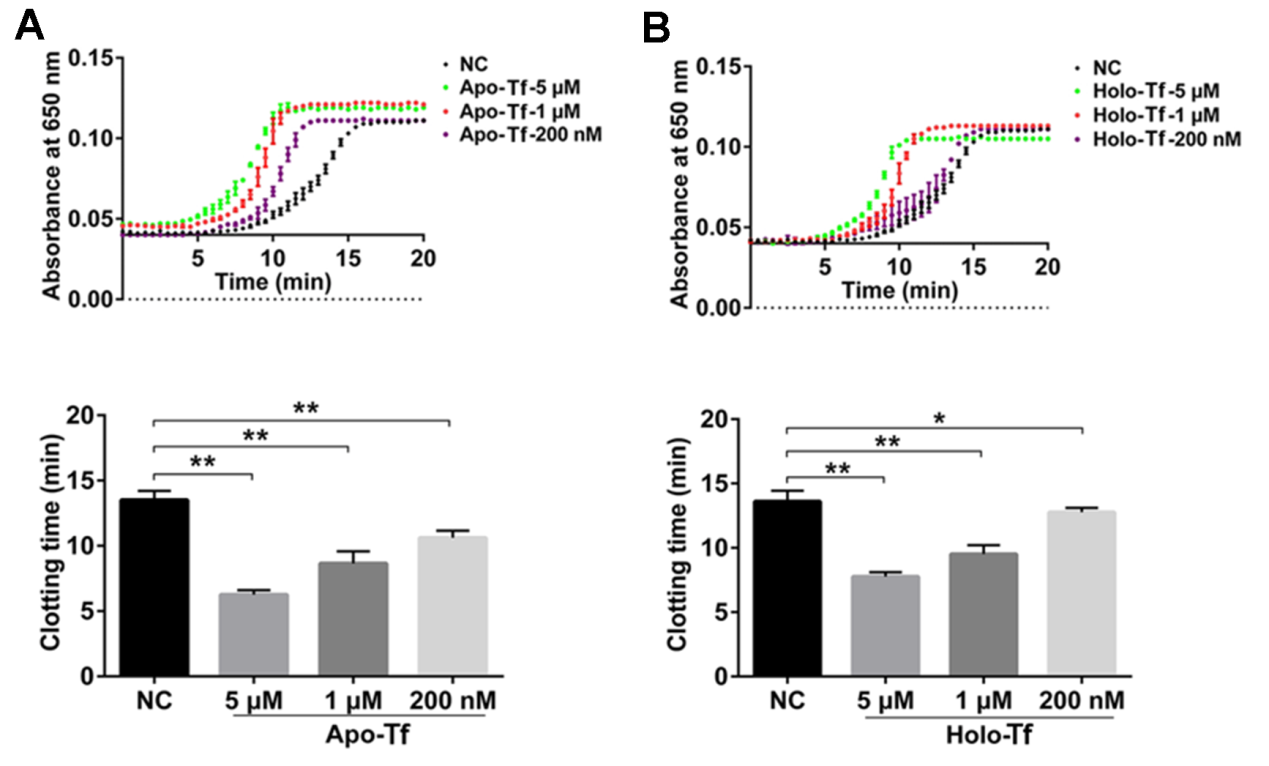

**Figure S4. The effects of apo- and holo-Tf on plasma recalcification time.** Human plasma recalcification time was shortened by human apo- **(A)** or holo-Tf **(B)**. Clotting time (below) was calculated by measuring the time to half maximal increase in absorbance. Data represent mean ± SD of six independent experiments, **p* < 0.05, ***p* < 0.01 by one-way ANOVA with Dunnett’s post hoc test.

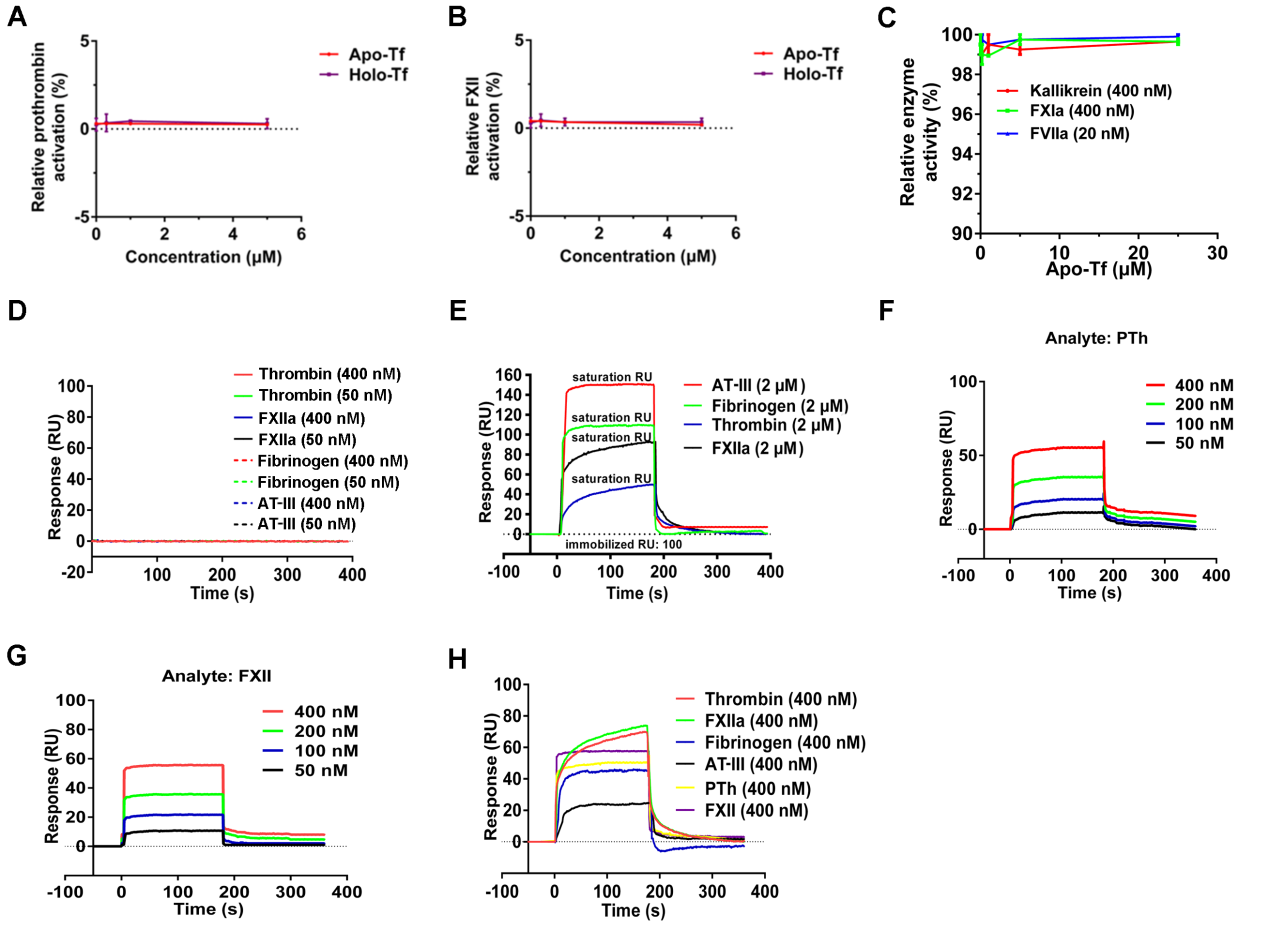

**Figure S5. The effects of apo- and holo-Tf on prothrombin/FXII activation and SPR analysis of the interaction between Tf and prothrombin/FXII.** Both apo- and holo-Tf showed no influence on prothrombin **(A)** or FXII **(B)** activation. Data represent mean ± SD of four independent experiments. **(C)** Apo-Tf showed no influence on kallikrein, FXIa and FVIIa. Data represent mean ± SD of six independent experiments**.** **(D)** SPR analysis of the interactions between human serum albumin (HSA) and thrombin, FXIIa, fibrinogen or antithrombin III (AT-III). **(E)** SPR analysis of binding molar ratio between Tf and fibrinogen, AT-III, thrombin and FXIIa. SPR analysis of the interaction between Tf and prothrombin **(F)** and FXII **(G)**. **(H)** SPR analysis of the interactions between holo-Tf and thrombin, FXIIa, fibrinogen, AT-III, prothrombin (PTh) and FXII.

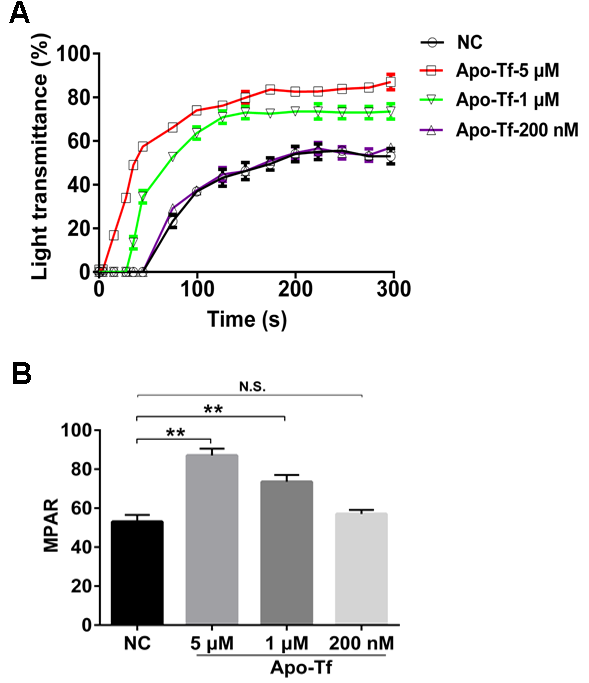
**Figure S6. The effect of apo-Tf on thrombin-induced platelet aggregation.** Tf’s effect on thrombin-induced platelet aggregation **(A)** and the maximum platelet aggregation rate (MPAR) **(B)**. Data represent mean ± SD of six independent experiments, ***p* < 0.01 by one-way ANOVA with Dunnett’s post hoc test.

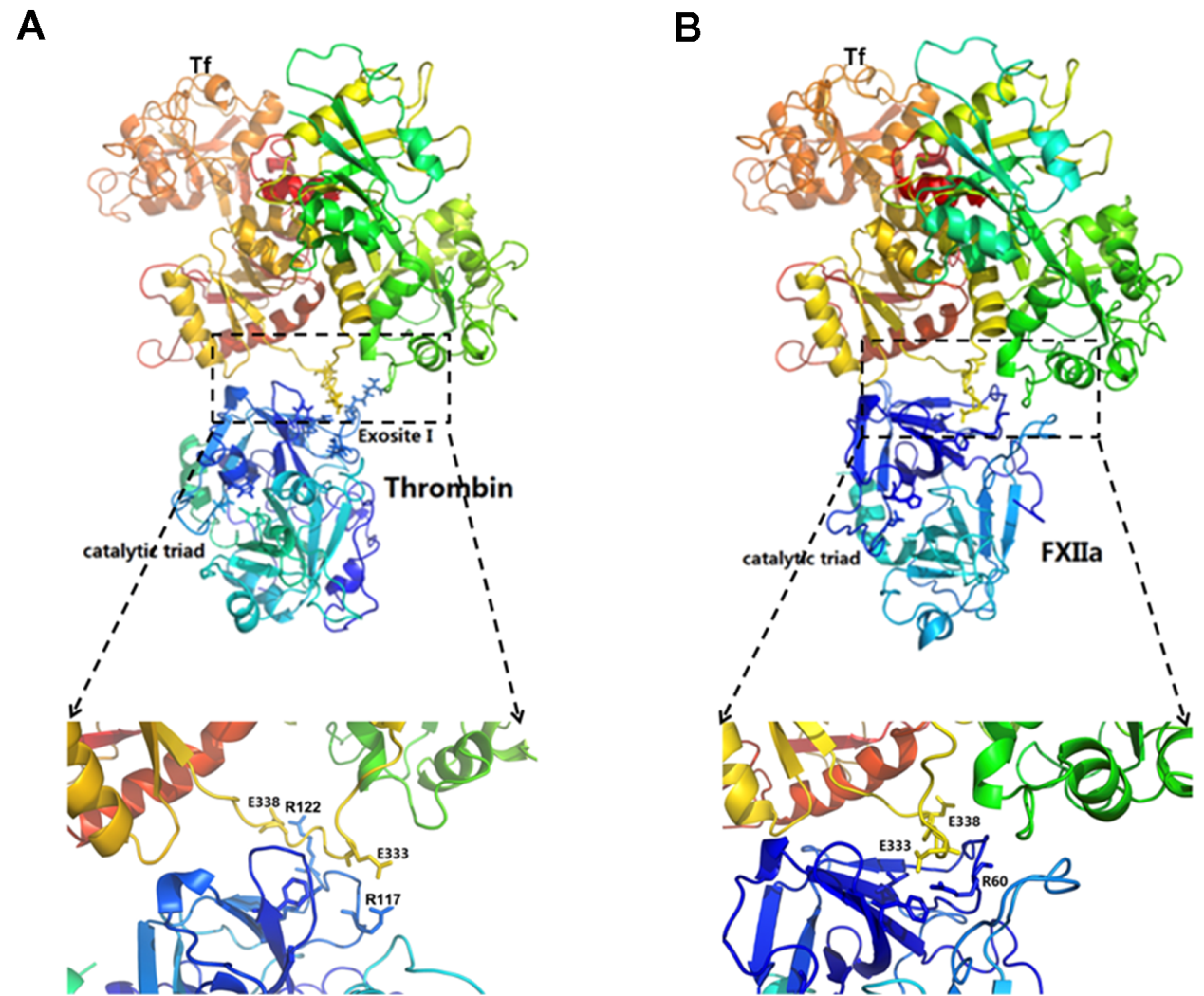

**Figure S7.** **Interaction of Tf with thrombin and FXIIa by molecular docking. (A)** Tf located in the exosite I of human thrombin. The homological model of human thrombin was constructed from the known structures (PDB ID: 4NZQ) and indicated by the illustration with key residues highlighted by a stick representation. The catalytic triad was marked by colorful sticks of side chains. **(B)** Two structures were used to build the working model of human FXIIa (PDB ID 4XE4 and 4XDE). This FXIIa model was aligned along the thrombin model to confirm the possible groove-like binding area.

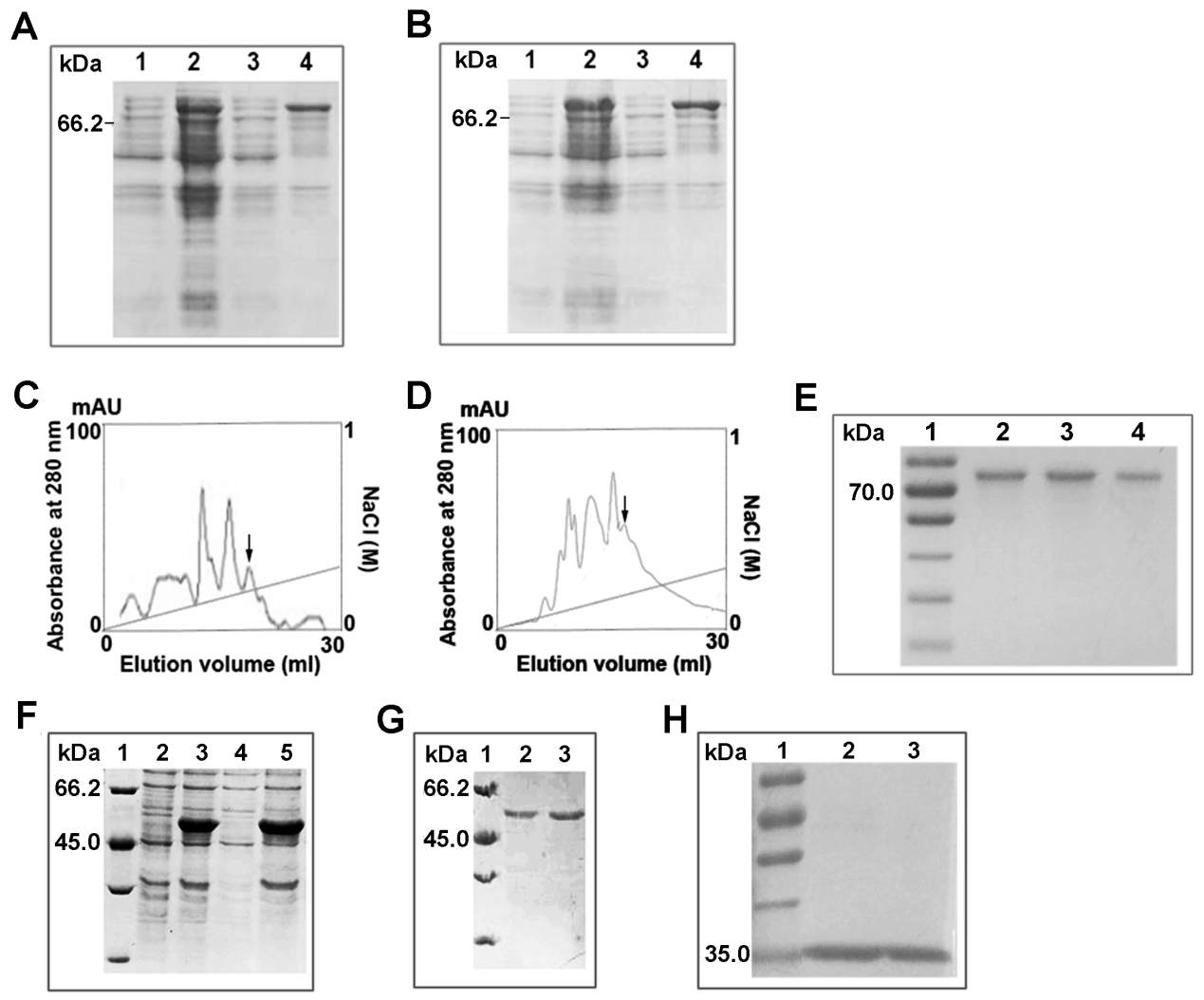

**Figure S8. Recombinant expression and purification of Tf, thrombin and their mutants. (A)** SDS-PAGE (12%) analysis of the production of fused wild-type Tf after IPTG induction. Lane 1: non-induced; Lane 2: induced; Lane 3: fraction containing wild-type Tf fusion protein eluted from Ni^2+^ affinity chromatography column was cut by enterokinase (5 unit); Lane 4: fraction containing wild-type Tf fusion protein eluted from Ni^2+^ affinity chromatography column. **(B)** SDS-PAGE (12%) analysis of the production of fused Tf mutant after IPTG induction. Lane 1: non-induced; Lane 2: induced; Lane 3: fraction containing Tf mutant fusion protein eluted from Ni^2+^ affinity chromatography column was cut by enterokinase (5 unit); Lane 4: fraction containing Tf mutant fusion protein eluted from Ni^2+^ affinity chromatography column. The fraction containing wild-type Tf **(C)** and Tf mutant **(D)** was eluted by Mono Q column, respectively. The gray line represents a line gradient of solvent B (20 mM Tris-HCl, 1 M NaCl, pH 7.8) from 0 to 30% over 30 min. Arrows indicate purified wild-type Tf and Tf mutant. **(E)** SDS-PAGE analysis of purified wild-type Tf and Tf mutant in a 12% gel. Lane 1: protein markers; Lane 2: apo-Tf (T2036, Sigma); Lane 3: purified wild-type Tf; Lane 4: purified Tf mutant. **(F)** SDS-PAGE analysis of the production of fused wild-type thrombin and thrombin mutant after IPTG induction. Lane 1: protein marker; Lane 2: wild-type thrombin, non-induced; Lane 3: wild-type thrombin, induced; Lane 4: thrombin mutant, non-induced; Lane 5: thrombin mutant, induced. **(G)** The bound fusion protein of Ni^2+^ affinity chromatography column; Lane1: protein marker; Lane2: fusion protein of wild-type thrombin; Lane3: fusion protein of thrombin mutant. **(H)** Purified wild-type thrombin or thrombin mutant. Lane 1: protein marker; Lane 2: purified wild-type thrombin; Lane 3: purified thrombin mutant.

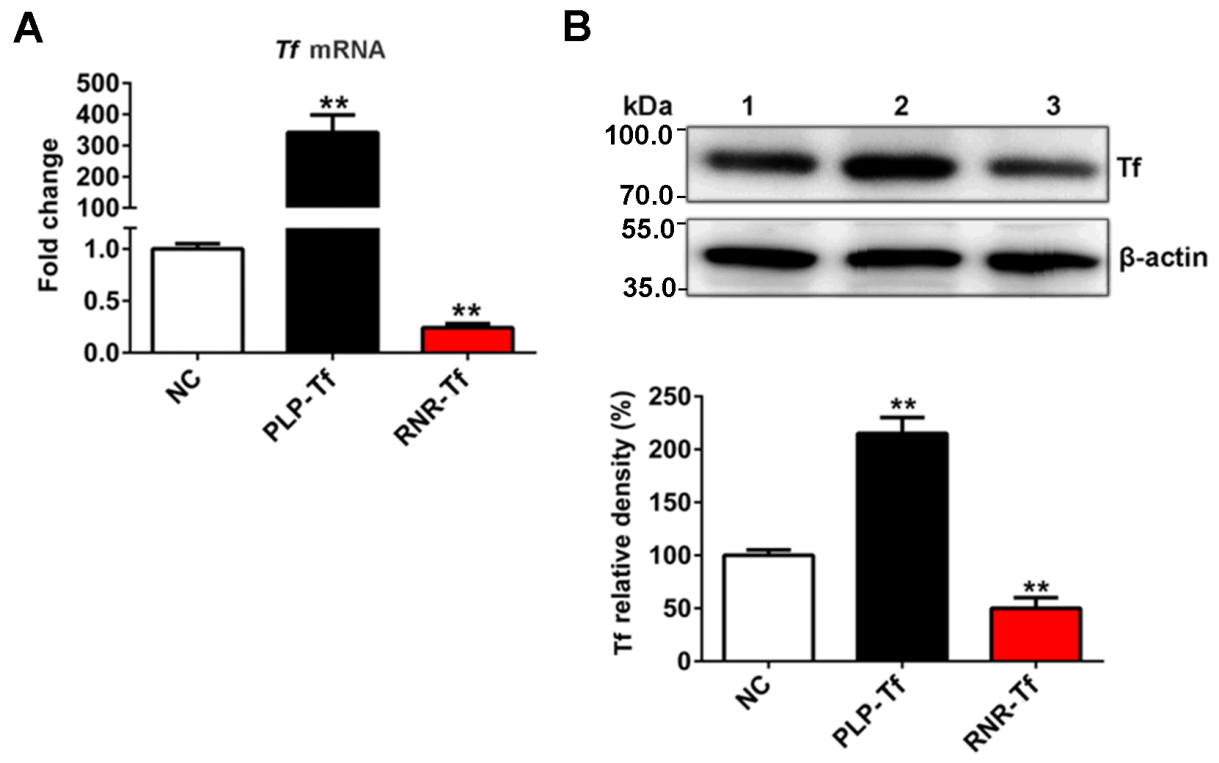

**Figure S9.** **Construction of Tf over-expression and knockdown vectors.** **(A)** Tf mRNA levels of BNL CL.2 cells after transfecting the over-expression or knockdown plasmid of Tf were determined by qRT-PCR. **(B)** Tf levels in the BNL CL.2 cells were determined by western blot (top, Lane 1: control (NC), Lane 2: over-expression (PLP-Tf), Lane 3: knockdown (RNR-Tf)). The quantification of the result of western blot was also shown (bottom). Data represent mean ± SD of six independent experiments, ***p* < 0.01 by one-way ANOVA with Dunnett’s post hoc test.

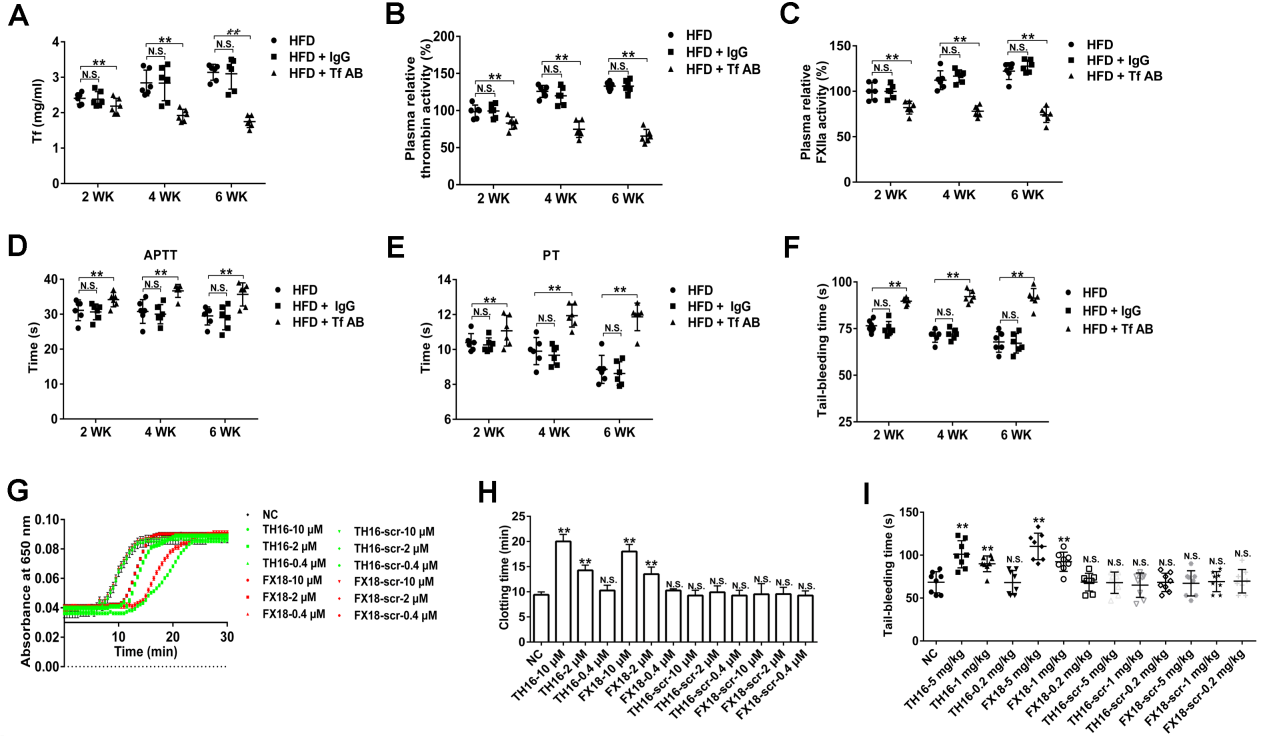

**Figure S10. Tf interferences inhibit hypercoagulability *in vivo*.** The HFD-fed *Apoe^-/-^* mice were subjected to anti-Tf antibody (Tf AB) or IgG treatment 2 times/week for 6 weeks. Plasma concentrations of Tf **(A)**, relative thrombin **(B)** and FXIIa activities **(C)** in plasma, APTT **(D)**, PT **(E)** and tail-bleeding time **(F)** in *Apoe^-/-^* mice were shown. Effects of TH16, FX18, TH16-scr and FX18-scr on plasma recalcification time **(G)**, clotting time **(H)** and tail-bleeding time **(I)** in C57BL/6J mice. Data represent mean ± SD (n = 6-8), **p < 0.01 by one-way ANOVA with Dunnett’s post hoc test. N.S.: no significance.

**Supplementary Tables**

**Table S1. Clinical features and plasma Tf concentrations of patients with CHD.**

| Sex | Age (years) | Main clinical features | Tf (mg/ml) |
| --- | --- | --- | --- |
| CHD patients | | | |
| Female | 54 | AP, AVLN | 5.300 |
| Female | 55 | AP, AVLN | 3.000 |
| Female | 61 | AP, AVLN | 3.700 |
| Male | 52 | AP, AVLN | 5.900 |
| Female | 68 | AP, AVLN | 5.136 |
| Female | 71 | AP, AVLN | 4.200 |
| Male | 64 | AP, AVLN | 2.976 |
| Female | 63 | AP, AVLN | 5.136 |
| Male | 43 | AP, AVLN | 2.900 |
| Male | 59 | AP, AVLN | 7.080 |
| Female | 68 | AP, AVLN | 4.104 |
| Male | 74 | AP, AVLN | 4.600 |
| Male | 60 | AP, AVLN | 3.312 |
| Male | 49 | AP, AVLN | 3.312 |
| Male | 57 | AP, AVLN | 4.104 |
| Male | 35 | AP, AVLN | 3.200 |
| Female | 69 | AP, AVLN | 3.096 |
| Male | 70 | AP, AVLN | 2.904 |
| Female | 44 | AP, AVLN | 3.312 |
| Female | 65 | AP, AVLN | 7.164 |
| Female | 79 | AP, AVLN | 6.600 |
| Male | 57 | AP, AVLN | 3.600 |
| Female | 39 | AP, AVLN | 4.980 |
| Female | 71 | AP, AVLN | 3.156 |
| Female | 67 | AP, AVLN | 3.228 |
| Female | 57 | AP, AVLN | 2.990 |
| Male | 61 | AP, AVLN | 3.312 |
| Female | 58 | AP, AVLN | 5.500 |
| Male | 64 | AP, AVLN | 3.312 |
| Male | 70 | AP, AVLN | 3.336 |
| Male | 61 | AP, AVLN | 3.312 |
| Female | 66 | AP, AVLN | 3.500 |
| Female | 71 | AP, AVLN | 5.136 |
| Female | 38 | AP, AVLN | 3.200 |
| Male | 63 | AP, AVLN | 3.976 |
| Male | 29 | AP, AVLN | 6.720 |
| Female | 46 | AP, AVLN | 5.148 |
| Male | 58 | AP, AVLN | 3.312 |
| Male | 39 | AP, AVLN | 3.336 |
| Male | 71 | AP, AVLN | 5.148 |
| Male | 56 | AP, AVLN | 3.120 |
| Male | 67 | AP, AVLN | 4.900 |
| Controls (healthy volunteers) | | | |
| Male | 39 | NA | 2.59 |
| Male | 48 | NA | 2.55 |
| Male | 34 | NA | 3.60 |
| Male | 54 | NA | 2.47 |
| Female | 38 | NA | 2.59 |
| Female | 27 | NA | 3.02 |
| Female | 60 | NA | 2.59 |
| Female | 27 | NA | 2.54 |
| Male | 65 | NA | 2.80 |
| Male | 33 | NA | 2.54 |
| Female | 43 | NA | 2.80 |
| Male | 59 | NA | 3.24 |
| Female | 39 | NA | 2.82 |
| Female | 51 | NA | 3.35 |
| Male | 67 | NA | 3.53 |
| Female | 51 | NA | 3.43 |
| Male | 45 | NA | 3.24 |
| Female | 36 | NA | 3.20 |
| Female | 39 | NA | 3.27 |
| Male | 47 | NA | 2.80 |
| Male | 29 | NA | 2.80 |
| Male | 42 | NA | 3.53 |
| Male | 67 | NA | 2.67 |
| Male | 45 | NA | 3.13 |
| Female | 42 | NA | 2.85 |
| Female | 65 | NA | 2.82 |
| Female | 27 | NA | 2.82 |
| Male | 45 | NA | 3.27 |
| Male | 45 | NA | 2.57 |
| Male | 62 | NA | 3.21 |
| Female | 33 | NA | 3.20 |
| Male | 45 | NA | 2.72 |
| Female | 27 | NA | 2.71 |
| Female | 37 | NA | 2.93 |
| Female | 34 | NA | 2.90 |
| Female | 76 | NA | 2.58 |
| Male | 57 | NA | 2.42 |
| Female | 21 | NA | 2.58 |
| Female | 49 | NA | 2.51 |
| Male | 36 | NA | 2.14 |
| Female | 32 | NA | 2.40 |
| Male | 38 | NA | 2.60 |

AP: angina pectoris; AVLN: angiographically visible luminal narrowing; NA: not applicable.

**Table S2. Clinical features of atherosclerotic plaque donors.**

| Sex | Age (years) | Main clinical features |
| --- | --- | --- |
| AS plaque donors | | |
| Male | 70 | AP, AVLN |
| Male | 67 | AP, AVLN |
| Male | 65 | AP, AVLN |
| Female | 49 | AP, AVLN |
| Female | 54 | AP, AVLN |
| Female | 60 | AP, AVLN |
| Female | 63 | AP, AVLN |
| Male | 42 | AP, AVLN |
| Female | 50 | AP, AVLN |
| Female | 68 | AP, AVLN |
| Male | 71 | AP, AVLN |
| Male | 63 | AP, AVLN |

AP: angina pectoris; AVLN: angiographically visible luminal narrowing.

**Table S3. Iron metabolism indexes (iron, ferritin) and erythrocyte indexes (MCV, MCH, MCHC) of normal mice (NC), Tf over expression (PLP-Tf), knock down (RNR-Tf), anti-Tf antibody (Tf AB) and IgG treatment mice.**

|  | Iron  (μM/L) | Ferritin (μg/L) | MCV  (fL) | MCH  (pg) | MCHC (g/L) |
| --- | --- | --- | --- | --- | --- |
| NC | 16.4 (1.34) | 72.2 (2.97) | 44.8 (2.12) | 14.2 (1.21) | 307 (6.73) |
| PLP-Tf | 16.1 (1.96) | 71.5 (3.45) | 43.5 (3.91) | 13.9 (1.98) | 310 (5.89) |
| RNR-Tf | 16.5 (1.15) | 70.9 (1.95) | 44.1 (2.54) | 14.4 (1.31) | 309 (7.21) |
| Tf AB | 16.0 (1.52) | 73 (3.14) | 44.0 (2.34) | 14.5 (1.34) | 312 (6.21) |
| IgG | 16.2 (1.23) | 71 (2.34) | 45.1 (3.24) | 14.1 (2.11) | 305 (5.89) |

MCV: mean corpusular volume; MCH: mean corpusular hemoglobin; MCHC: mean corpuscular hemoglobin concentration. Data represent mean (SD), n = 10.
